# Characterization of Orsay virus replication intermediates in *Caenorhabditis elegans* reveals links to antiviral RNA interference

**DOI:** 10.1101/2025.09.15.676353

**Authors:** Supraja Ranganathan, P. Joseph Aruscavage, Brenda L. Bass

**Affiliations:** Department of Biochemistry, University of Utah, Salt Lake City, Utah, USA 84112-5650

## Abstract

Orsay Virus (OV) is a positive-sense, single-stranded RNA (+ssRNA) virus that naturally infects *C. elegans* intestines. Like other +ssRNA viruses, the OV-encoded RNA-dependent RNA polymerase (oRdRP) synthesizes complementary antigenome for use as template for amplifying viral genome, but OV replication intermediates are underexplored. Using PCR, we observed viral genome in vast excess of antigenome, as for other +ssRNA viruses. Unlike interferon-based antiviral defense, *C. elegans* utilizes RNA interference (RNAi) for antiviral defense, producing sense and antisense small interfering RNAs (siRNAs) that cannot be distinguished from genome and antigenome with conventional hybridization protocols. Fluorescence-based imaging in *C. elegans* intestines using probes to antigenomic sequences revealed cytoplasmic as well as perinuclear localization patterns. The latter depended on factors required for generation of primary, but not secondary, siRNAs, connecting the antigenomic hybridization pattern to RNAi. We also observed cytoplasmic double-stranded RNA (dsRNA) associated with oRdRP, suggesting viral replication hubs, as well as infection-induced nuclear dsRNA, likely from endogenous dsRNA. Finally, using antibodies to oRdRP, we observed spherical structures of ∼1µm in diameter with oRdRP at their surface, which decrease in animals lacking RDE-1. Our study defines features of OV replication intermediates, setting the stage for understanding their connection to host antiviral pathways.

**Significance Statement:**

- Orsay virus is a +ssRNA virus that infects *C. elegans* intestines. We advance understanding of viral replication intermediates and address the issue that for animals that use antiviral RNA interference, hybridization of probes occurs with both genome and antigenome and small interfering RNAs.
- Single-molecule fluorescence in-situ hybridization using antigenomic probes revealed cytoplasmic and perinuclear puncta, only upon denaturation, with perinuclear signal dependent on primary, but not secondary, siRNA biogenesis. Viral RNA-dependent RNA polymerase lined the perimeter of spherical structures of ∼1µm diameter.
- This study sets the stage for understanding the relationship between viral replication dynamics and antiviral RNA interference.

## Introduction

Orsay Virus (OV) is a naturally infecting *C. elegans* pathogen that closely resembles the *Nodaviridae* family of viruses. OV is a positive strand, single-stranded RNA (+ssRNA) virus, with a bipartite genome (oRNA1 and oRNA2), that preferentially infects *C. elegans* intestinal cells where it is predominantly restricted to anterior cells (Félix *et al*., 2011; Franz *et al*., 2014; Castiglioni *et al*., 2024). oRNA1 encodes an RNA-dependent RNA Polymerase (oRdRP), and oRNA2 encodes the capsid, δ (delta), and capsid-δ fusion proteins (Félix and Wang, 2019).

While details of OV replication and its intermediates are just emerging (Franz *et al*., 2014; Batachari *et al*., 2024), it can be presumed that general features of positive-strand RNA viruses are operative (de Beijer *et al*., 2024; Stancheva and Sanyal, 2024). Replication of a positive-strand RNA virus begins with translation of its RdRP from the genomic strand. The RdRP first synthesizes a complementary strand to the genome, the antigenome (-ssRNA), and this is the first RNA replication intermediate. The antigenome is then used as a template by the RdRP to amplify the viral genome (Ball, 1994). The perfectly complementary strands of genome and antigenome can basepair to form a viral double-stranded RNA (dsRNA), another RNA intermediate of viral replication. Replication of closely related nodaviruses occurs inside membrane-enclosed compartments that protect viral RNA and other replication intermediates from the host antiviral machinery (Stancheva and Sanyal, 2024). The structural proteins along with the viral genome they encapsulate are packaged and processed into a mature virion that is then egressed out of the cell.

Some viral replication intermediates are recognized by helicases to elicit an antiviral response. Among vertebrates, the RIG-I-like Receptor (RLR) family of helicases recognize dsRNA to elicit an interferon response. Invertebrates do not have interferon pathways, and invertebrates such as *C. elegans* instead utilize RNA interference (RNAi). In RNAi it is the enzyme Dicer that recognizes dsRNA, using a helicase domain related to that of RLRs. *C. elegans* Dicer (DCR-1) forms an antiviral RNAi complex with DRH-1, a Dicer-related helicase and RLR ortholog, and RDE-4 (RNAi-DEficient 4), a dsRNA binding protein. This antiviral complex (AVC) cleaves viral dsRNA into 23H (H=A,U,C>G at 5’ end) primary small-interfering RNAs (1° siRNAs). The 1° siRNA is then bound by RDE-1 (RNAi-deficient 1), an argonaute protein, and passed to other factors for generation and amplification of 22G (preferentially G at 5’ end) secondary siRNAs (2° siRNAs) that are complementary to the viral genome (Ashe *et al*., 2013; Guo *et al*., 2013). In the germline, amplification of 2° siRNAs occurs in perinuclear compartments that are marked by the presence of RNAi amplification factors such as RRF-1 and DRH-3 (Phillips *et al*., 2012). However, very little is known about the spatial organization of RNAi factors and RNAi hubs in somatic tissues, such as the intestine, and during OV infection.

Somewhat analogous to the mammalian interferon response, the *C. elegans* antiviral response also involves transcription of OV-induced genes (OVIGs) (Chen *et al*., 2017; Castiglioni *et al*., 2024), a subset of which are the *C. elegans* Intracellular Pathogen Response (IPR) genes (Reddy *et al*., 2017); transcription of OVIGs requires DRH-1, but not DCR-1 or RDE-4 (Sowa *et al*., 2019). DRH-1-mediated OVIG induction occurs after viral replication, but the exact viral intermediate that DRH-1 recognizes to trigger OVIGs is unknown. However, recently OV dsRNA was visualized in association with DRH-1 in *C. elegans* intestines, raising the possibility that this recognition is important for triggering OVIGs (Batachari *et al*., 2024). *C. elegans* lacking RNAi factors have increased sensitivity to OV, and these mutant strains are widely used to understand OV replication dynamics and downstream RNAi responses (Franz *et al*., 2014). Given that replication is required to elicit host antiviral responses, it is important to understand the temporal and spatial organization of replication intermediates. The most well-studied viral replication byproduct is viral dsRNA. We know less about other replication intermediates, leading to an incomplete picture of replication dynamics and antiviral responses, and for *C. elegans*, this is especially true for a wildtype background.

Here we describe studies designed to detect and characterize three replication intermediates, oRNA1 antigenome, OV dsRNA, and oRdRP, in both wildtype and RNAi mutant backgrounds. We show that OV is like other +ssRNA viruses in that the viral genome is present in vast abundance relative to the antigenome, in both wildtype and RNAi mutants. Single molecule FISH (smFISH) using probes designed to detect antigenomic sequences show signal in cells with genomic strand, and detection is facilitated with denaturation protocols, indicating most antigenomic sequences are basepaired. Apart from the expected cytoplasmic distribution, unexpectedly, we observed signal with antigenomic probes within perinuclear regions; fewer animals showed this signal in strains defective for primary siRNA biogenesis, but not those aberrant for secondary siRNA amplification, setting the stage for connecting OV infection and replication with the antiviral RNAi pathway. We did not detect RNAi factors associating with the perinuclear-localized antigenomic signal, unlike *C. elegans* germline. We observed oRdRP-associated dsRNA puncta revealing possible replication hubs. Infected wildtype intestinal cells also showed oRdRP in lumen-containing spherical structures of ∼1μm in diameter that were dependent on factors of the antiviral RNAi pathway. Together, the results reveal new hubs of viral replication and processing, specifically in a wildtype background.

## Results

### OV genomic strand accumulates significantly more than antigenomic strand in wildtype and RNAi mutants

OV infection in *C. elegans* is typically measured by RT-qPCR in a strand-agnostic manner and denoted as viral load, or by monitoring genomic strand accumulation using northern blot analysis. For a plus-strand virus such as OV, these methods largely reflect levels of the more abundant genome but offer little information about the less abundant antigenome (Félix *et al*., 2011). To acquire this information, we first monitored OV-infected animals with strand-specific RT-qPCR. We designed strand-specific probes for oRNA1 genome or antigenome that each included a unique flanking sequence so that strand-specific complementary DNA (cDNA) could be quantified with qPCR, using the unique flanking region as a primer and a second downstream on the respective oRNA1 strand (**Figure 1A**). We also performed northern blot analysis to complement our strand-specific RT-qPCR method and directly detect viral RNA abundance.

**Figure 1:**
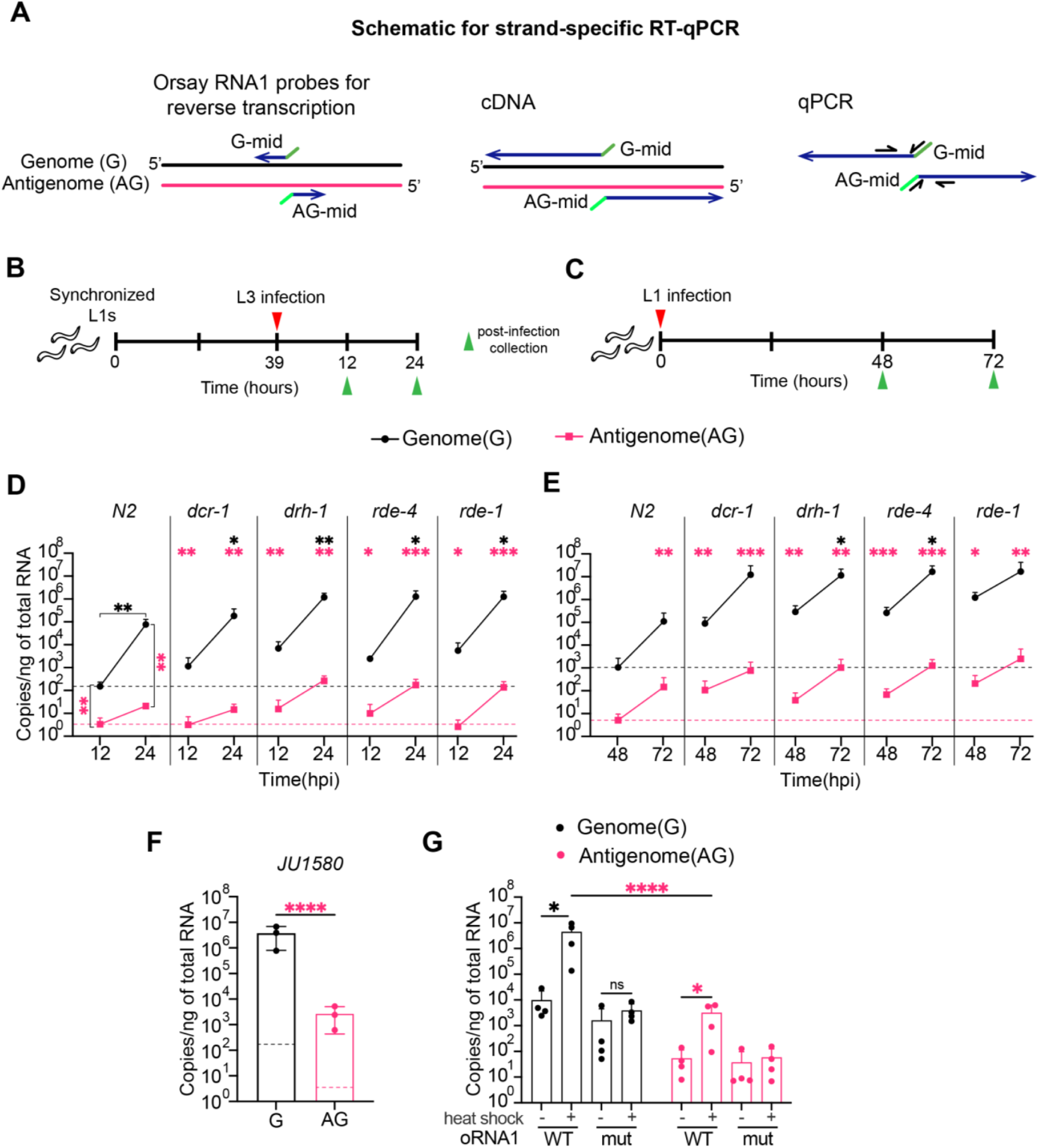
Dynamics of oRNA1 strand-specific accumulation upon OV infection. (A) Schematic of strand-specific RT-qPCR of OV RNA1. Primers with unique adapters (green colors) were used for reverse transcription to generate cDNA specifically from oRNA1 strands. Resulting cDNA was used for qPCR with primers that recognize the unique adapter and a downstream region (in black, half arrows). Schematics of (B) late larval (L3) and (C) early larval (L1) OV infection protocols. Number of copies (per ng of total RNA) of oRNA1 genome and antigenome at (D) L3 or (E) L1 infection of indicated strains and times. Data, mean with standard deviation (N=3, biological replicates). Black asterisks, significance between indicated infection time point and previous time point for oRNA1 genomic strand; Two-way ANOVA, Uncorrected Fischer’s LSD test. Pink asterisks, significance between genome and antigenome at indicated timepoint as shown for N2; Ratio paired t-test. No asterisks, no significant difference (See also S1 Fig). Number of copies of oRNA1 genome or antigenome as measured by strand-specific RT-qPCR in (F) constitutively OV-infected JU1580 animals (Dotted lines, copy number of genome or antigenome in N2 animals at 12hpi. Ratio paired t-test), and (G) in strains expressing oRNA1^WT^ or oRNA1^mut^ with or without heat shock. Data are mean with standard deviation (N=4, biological replicates). Ratio paired t-test. *\*, p-value<0.05; **, p-value<0.01; ***, p-value<0.001; ****, p-value<0.0001*.

We in vitro transcribed and purified fragments of genomic and antigenomic strands of oRNA1, each ∼500 nucleotides (nts) long, and used these transcripts to generate standard curves for quantifying copy number in a strand-specific manner (**Supplemental Figures 1A and 1B**); these transcripts were also used to confirm that antigenomic probes were specific and able to detect 10^6^ copies of purified ∼500nt antigenome ssRNA by northern blot assay (**Supplemental Figure 1C)**. We then monitored appearance of genome and antigenome over time, using two infection protocols: one monitoring infection of L3 larvae at 12- and 24-hours post infection (hpi; **Figure 1B**), and a second aimed at quantifying strands after prolonged infection, by infecting L1 larvae and assaying 48 and 72hpi (**Figure 1C**). For the latter, we chose infection times that maintained animals in the same reproductive cycle (72hpi) so that populations would not be contaminated with newly hatched larvae.

For each protocol, we compared infection of wildtype (N2) animals to that observed in antiviral RNAi mutants, *drh-1(uu60), rde-4(uu71), rde-1(ne219)*, and a helicase point mutant (G492R) of DCR-1 (*eri-4(mg375)*) (Pavelec *et al*., 2009a; Welker *et al*., 2010), referred to here as *dcr-1(mg375)* (**Figures 1D and 1E**). We observed significantly higher levels of genome than antigenome for wildtype and RNAi mutant strains. Further, for each infection protocol, with few exceptions, we observed a dramatic and significant increase in oRNA1 genome copy numbers at 24hpi and 72hpi relative to 12hpi or 48hpi, respectively, in both wildtype and mutant animals. We observed a modest increase in oRNA1 antigenome relative to wildtype in *drh-1, rde-4* or *rde-1* mutants, but at early time points this was not observed in *dcr-1(mg375)* mutants (**Figures 1D and 1E; points relative to pink dotted line**). Quantification showed greater than 100-fold increase in genome over antigenome at 12hpi, and over 1000-fold increase at all other infection times, for both wildtype and RNAi mutants (**Supplemental Figures 1D and 1E**). Northern blot analyses confirmed the low levels of antigenome compared to genome (**Supplemental Figures 1F and 1G**); controls established that if present antigenome could be detected, but oRNA1 antigenome was below the limit of detection for wildtype and RNAi mutants at all time points tested (12-72 hpi). oRNA1 genome was first detected at 24hpi for RNAi mutants, and increased over time, but was only faintly detected at 72hpi for wildtype strains.

Consistently, levels of genomic RNA were higher in RNAi mutants than wildtype (**Figures 1D and 1E**), and trends matched the increase in viral load as determined in a strand-agnostic manner, consistent with prior studies (Ashe *et al*., 2013; Sowa *et al*., 2019) (**Supplemental Figures 1H and 1I).** In accordance with published studies, we observed that the antiviral RNAi mutants, *drh-1*, *rde-4* and *rde-1*, were sensitive to OV and showed increased viral load relative to wildtype at all timepoints tested, by all methods (Ashe *et al*., 2013). While previous studies have reported the sensitivity of *dcr-1* using deletion mutants or knockdown (RNAi) (Ashe *et al*., 2013; Sowa *et al*., 2019), we report the involvement of the helicase domain of DCR-1. The helicase mutant *dcr-1(mg375)* showed a delayed response with a moderate increase in viral load, ∼10-fold at 12hpi and 24hpi, and ∼100-fold at 48hpi and 72hpi **(Figures 1D and 1E; Supplemental Figures 1H and 1I).**

Prior studies have looked at viral infection in the constitutively infected JU1580 strain, which has a mutation in DRH-1 and is highly sensitive to OV infection (Félix *et al*., 2011; Félix and Wang, 2019). To connect our studies to prior studies we showed that a high viral load in the JU1580 strain relative to wildtype, as determined using strand-agnostic protocols, correlated with a robust accumulation of oRNA1 genomic strand by both strand-specific RT-qPCR and northern blot, while the oRNA1 antigenome levels remained significantly lower, ∼1300-fold, (*p-value<0.0001*) (**Figures 1F; Supplemental Figures 1F and 1G**). This emphasizes that the antigenomic strand is much lower than the genomic strand even with constitutive infection. To test if the ratio of genome to antigenome was a property inherent to oRdRP mediated replication, we also monitored a heat shock-inducible replicon system expressing wildtype (oRdRP^WT^) or a replication-incompetent mutant (oRdRP^mut^) oRNA1 (Jiang *et al*., 2017). In the complete absence of infection, we observed a significant increase in genome and antigenome upon inducing the expression of oRdRP^WT^ but not oRdRP^mut^, and notably, antigenome levels remained significantly lower (**Figure 1G**), which was confirmed by northern blot (**Supplemental Figure 1F**).

In summary, these studies indicate that the increase in viral load commonly measured using strand-agnostic methods largely reflects an increase in genomic strand. At the time points tested, for wildtype and RNAi mutant strains, we found that the genomic strand, the strand that is packaged into virions, is in far excess of the antigenomic strand, which is only an intermediate, and these ratios are intrinsic to the oRdRP and its tight regulation by host factors (Ahlquist *et al*., 2003; Jakubiec and Jupin, 2007).

### Signal using probes to the oRNA1 antigenome is present upon denaturation

To gain insight into the localization of OV genome and antigenome within infected cells, we performed smFISH analyses by designing multiple strand-specific fluorescent deoxyoligonucleotide probes for oRNA1 (27 probes, genomic strand; 48 probes, antigenomic strand). For initial optimization, we used OV infection-sensitive *jyIs8;rde-1* animals. *jyIs8* contains an integrated transgene that expresses GFP under the control of the promoter of an OVIG, *pals-5*, and was used as a marker for infection (Bakowski *et al*., 2014). Uninfected controls established the absence of signal for *pals-5p::GFP* and oRNA1 strands, while as in prior studies (Bakowski *et al*., 2014), a strong signal in the pharynx (asterisk) in the same channel as antigenome was observed from fluorescence of the *myo-2p::mCherry* marker included in the *jyIs8* transgene (**Figure 2A; uninfected**). At 24hpi we saw robust GFP expression in multiple intestinal cells that typically extended beyond the cell(s) that showed robust signal with probes to genomic sequences. The multi-cell GFP expression is consistent with prior studies showing distribution of DRH-1 and ZIP-1 beyond infected cells (Lažetić *et al*., 2022; Batachari *et al*., 2024). When we used antigenomic probes, ∼95% of infected animals showed a diffuse cytoplasmic signal without distinct puncta in the same intestinal cells that showed genomic signal (**Figures 2A, infected (-) denature and 2B**). Given the requisite complementarity between OV strands, we wondered if the observed signal was affected by basepairing. To test this, we treated animals with 50mM NaOH for ∼30 seconds to promote dsRNA denaturation prior to hybridization (Genoyer *et al*., 2025), hereafter referred to as denature-smFISH (dsmFISH). After denaturation, signal using genomic probes was unchanged, consistent with the conventional understanding of replication of a positive strand virus, whereby a vast excess of single-stranded genome is produced for subsequent packaging. By contrast, after denaturation, in addition to the diffuse cytoplasmic signal, antigenomic probes revealed a punctate signal in ∼97% of infected animals (**Figures 2A, infected (+) denature and 2B**). This indicated that the oRNA1 antigenomic signal derived from a species that was basepaired via intramolecular or intermolecular interactions. We used the dsmFISH denaturing conditions for subsequent experiments. Further, because the genomic signal was unaffected by denaturation, in subsequent experiments we interpreted signal from genomic probes as corresponding to the single-stranded genome quantified in our PCR experiments. The dependence of the antigenomic signal on denaturation suggested the signal derived from a basepaired species, either the lowly abundant antigenomic strand, viral dsRNA, or the viral siRNAs generated during antiviral RNAi.

**Figure 2:**
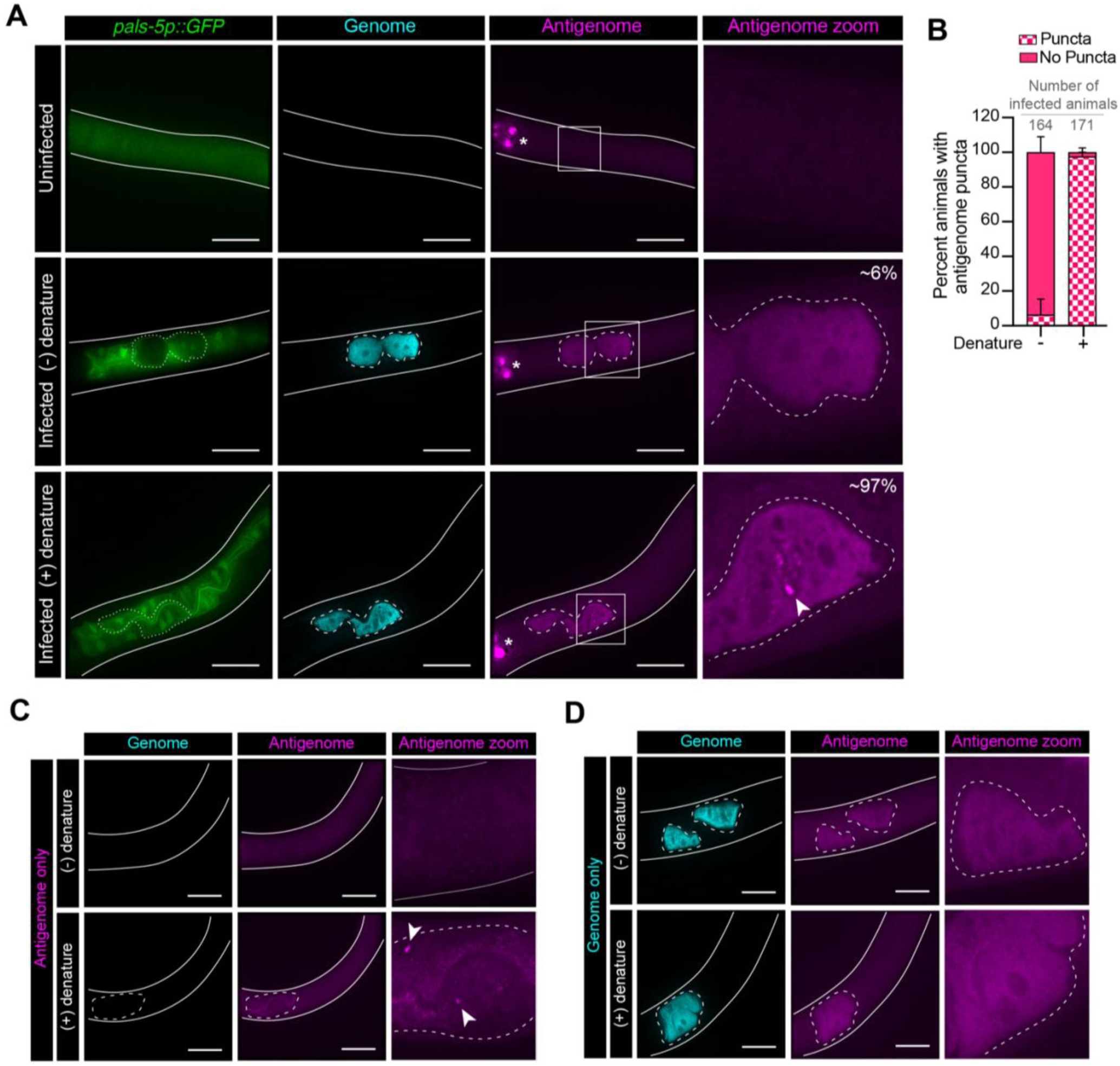
Signal from probes for antigenome is evident upon RNA denaturation. (A) Representative smFISH images of uninfected (denatured) and infected animals (*jyIs8;rde-1*) with or without dsRNA denaturation. *pals-5p::GFP* (green), oRNA1 genome (cyan), oRNA1 antigenome (magenta), scale bar, 25μm. Column 4 shows an enlarged image of region boxed in column 3. White arrowhead points to antigenome puncta. Pharynx signal (*) is from *mCherry*, a co-expression marker of the integrated transgene (*jyIs8*). (N>3, biological replicates). Solid white lines outline animals, dashed white lines outline infected intestinal cells. (B) Distribution of percent animals exhibiting antigenomic signal (puncta) with or without denaturation (N=3, biological replicates). Total number of infected animals is indicated. Representative images of OV-infected animals with or without denaturation using either (C) antigenome or (D) genome smFISH probes only. oRNA1 genome (cyan), oRNA1 antigenome (magenta), scale bar, 25μm. Solid white lines outline an animal, dashed white lines outline infected intestinal cells, and white arrowhead points to antigenome puncta in column 3 (antigenome zoom of C).

Since the diffuse cytoplasmic signal observed with probes to the antigenome was similar to the pattern of the oRNA1 genome signal, we wondered if the diffuse antigenomic signal was a consequence of bleed-though from the genome channel. Thus, we monitored antigenome in infected animals that had not been probed for genome, and vice-versa (**Figures 2C and 2D)**. We observed the diffuse signal in the antigenome channel even when we only probed for genome, consistent with bleed-through. In subsequent experiments we considered the diffuse cytoplasmic signal as nonspecific background. However, a lower level of cytoplasmic signal, and importantly, distinct puncta, were specific to antigenomic probes (**Figure 2C, white arrowhead**).

### Probes to the oRNA1 antigenome reveal perinuclear foci upon infection

Having established optimal protocols, we performed more extensive analyses, now comparing *jyIs8* (wildtype) with *jyIs8;rde-1* mutant animals at 24hpi. Again, infected intestinal cells showed robust genomic signal as well as distinct puncta when using antigenomic probes (**Figure 3A**). *rde-1* mutant animals exhibited more infected cells than wildtype as quantified by cytoplasmic genomic signal (**Figure 3B**). Further, antigenomic probes revealed a distinct perinuclear signal, emphasized by line scan analyses, in both wildtype and *rde-1* mutant strains (**Figure 3A)**; the perinuclear signal was only observed after denaturation and was also observed when probes to the antigenome were used exclusively, ruling out artifacts due to bleed-through (**Supplemental Figure 2**). Quantification showed that in wildtype or *rde-1* mutant strains, ∼83% or ∼78% of infected animals exhibited perinuclear signal, respectively (**Figure 3C**). The perinuclear puncta observed using antigenomic probes lacked an obvious colocalizing genomic signal, but the line scan showed a peak of closely associated genomic signal (**Figures 3D and 3E**). While the perinuclear puncta were the most common pattern observed with the antigenomic probes, multiple biological replicates (≥10) revealed other patterns in infected wildtype animals (**Figures 3F-J**). For example, close association of a punctate antigenomic signal with genomic signal was observed in the cytoplasm (**Figures 3F and 3G**). In other cases, antigenomic probes revealed concentrated cytoplasmic foci surrounded by genomic signal (**Figures 3H, white arrowhead and 3I**). Given that antigenomic strand is used as a template for viral replication to generate viral genome, possibly the closely associated genome and concentrated antigenomic puncta indicates active replication hubs in the infected cytoplasm. In infected animals that only exhibited genomic signal in the lumen **(Figure 3J**), as expected in animals that have released packaged virions (Castiglioni *et al*., 2024), we did not observe any cells with antigenomic signal, suggesting hybridizing material was degraded or processed through antiviral defense mechanisms, or below limits of detection.

**Figure 3:**
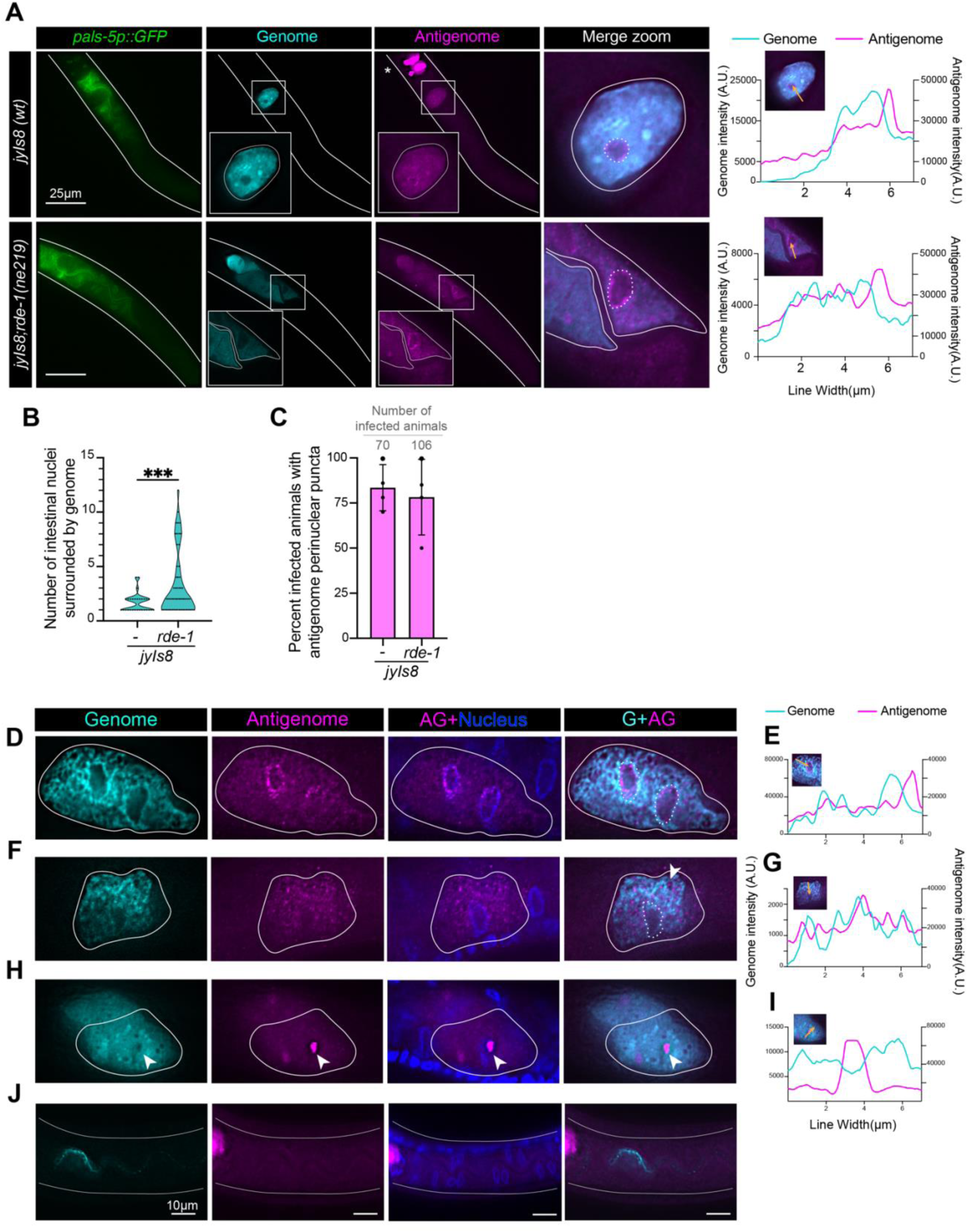
Probes to oRNA1 antigenome reveal perinuclear foci. (A) Representative images of *jyIs8(wt)* or *jyIs8;rde-1(ne219)* animals infected with OV for 24 hours along with corresponding line scan analysis (yellow arrow shows path of scan). Columns 1-3 depict *pals-5p::GFP* (green), oRNA1 genome (cyan), antigenome (magenta), respectively. Merge zoom (column 4) shows an enlarged image of the region boxed in columns 2 and 3 with signal from genome and antigenome. Scale bar, 25μm. Infected cells are outlined with a white solid line and nuclei (Hoechst) with a white dotted line. Pharynx signal (*) is from *mCherry,* a co-expression marker of the integrated transgene (*jyIs8*). (B) Violin plot showing number of nuclei surrounded by genome smFISH signal in indicated strains. Unpaired t-test with Welch’s correction, *\*\*\*; p=0.0003* (C) Bar graph showing percentage of infected animals exhibiting perinuclear localization of oRNA1 antigenome in infected cells of *jyIs8(wt)* or *jyIs8;rde-1(ne219).* N=4, biological replicates. Total number of infected animals is indicated in gray above each bar. (D-J) Representative images and line scans (yellow arrow shows path of scan) of *jyIs8 (wt)* infected cells (white solid line) showing distribution pattern of oRNA1 antigenomic signal (magenta) in (D, E) perinuclear, (F, G) cytoplasmic, or (H, I) cytoplasmic foci, in genome-containing infected cells, or (J) lumen (cyan, white solid line outlines infected animal). Dotted lines indicate nuclei in G+AG (column 4) to enable visualization of nuclei in other columns. In (H), showing cytoplasmic antigenome puncta, nucleus is unmarked since it is on a different z-plane.

### Primary siRNA biogenesis, but not secondary siRNA amplification, is required for perinuclear signal

A key aspect of antiviral defense in *C. elegans* is cleavage of viral dsRNA by the AVC, comprised of DCR-1, DRH-1, and RDE-4 (Consalvo *et al*., 2024). The AVC processes viral dsRNA to produce 23H double-stranded 1° siRNAs (Reich *et al*., 2018), and subsequently one strand of the 1° siRNA is bound by RDE-1 for presentation to factors that amplify the siRNA response by producing single-stranded 22G triphosphorylated 2° siRNAs. Mutations or deletions in DCR-1, DRH-1 or RDE-4 prevent accumulation of both 1° and 2° siRNAs, while RDE-1 mutants accumulate 1° siRNAs but lack 2° siRNAs (Ashe *et al*., 2013). In hopes of gaining insight into whether the perinuclear signal derived from full length antigenome or siRNAs, we infected synchronized L1 larvae of *dcr-1(mg375), drh-1(uu60), rde-4(uu71), rde-1(ne219)*, and wildtype (*N2*) animals for 24 hours and performed dsmFISH. As observed previously (**Figure 2A, uninfected**), a minimal background of diffuse fluorescence in the antigenomic channel in uninfected animals was observed for all strains (**Supplemental Figure 3**). At 24hpi, both wildtype and RNAi mutants showed abundant genome signal in one or more intestinal cells, with significantly more infected cells in RNAi mutants (**Figures 4A, G+AG column 1, and 4B**). Consistent with our previous experiments, we observed distinct perinuclear signal with antigenomic probes in wildtype (∼81% animals) or *rde-1* mutants (∼83% animals) (**Figures 4A and 4C, *N2, rde-1***). However, in animals mutant for one of the three factors of the AVC, a fraction of animals showed a loss of perinuclear signal (**Figures 4A and 4C, *dcr-1, drh-1, rde-4*)**. This result established a connection of the perinuclear signal to processing by the AVC, albeit does not definitively determine whether the perinuclear signal derives from the antigenome or antisense siRNAs. While animals with deficiencies in the AVC cannot process viral dsRNA as efficiently as wildtype, some activity remains, for example, certain DRH-1 mutations eliminate AVC processivity, but still generate siRNA reads that map to ends of the viral genome/antigenome (Ashe *et al*., 2013; Coffman *et al*., 2017). Possibly residual siRNAs allow a productive antiviral response in some but not all animals, explaining the reduction in animals with perinuclear puncta. MUT-16 is a scaffolding component required for mutator foci formation in the germline, sites where secondary siRNA amplification occurs (Phillips *et al*., 2012; Uebel *et al*., 2018). MUT-16 is also required for 2° siRNA amplification during OV infection (Lowe *et al*., 2025). We used a *mut-16(pk710)* strain to test if this strain phenocopies *rde-1* mutants with respect to perinuclear localization. We observed that over 85% of infected *mut-16* mutants at 24hpi exhibited perinuclear signal, not significantly different from wildtype or *rde-1* mutant backgrounds (**Figures 4A and 4C, *mut-16***). Since *rde-1* and *mut-16* mutant animals, which accumulate 1° siRNAs but lack 2° siRNAs, show robust perinuclear signal with antigenomic probes, our experiments suggest the perinuclear signal requires wildtype accumulation of 1° siRNAs but is unaffected by aberrant accumulation of 2° siRNAs.

**Figure 4:**
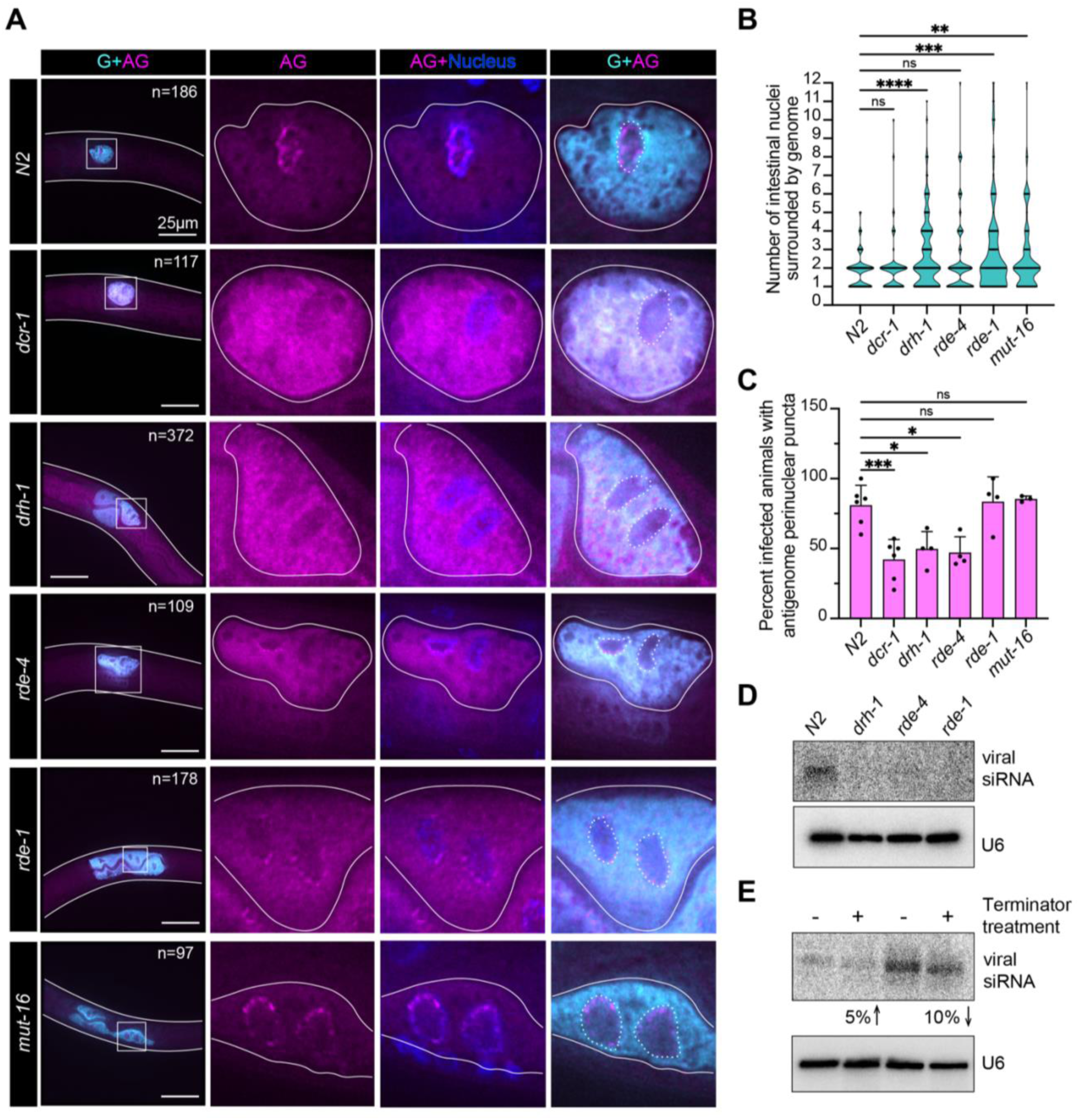
Perinuclear localization of oRNA1 antigenomic signal is partially dependent on primary siRNA biogenesis but not secondary siRNA amplification. (A) Representative images of OV-infected cells with distribution patterns of genome (G, cyan), antigenome (AG, magenta) and nuclei (Hoechst, blue) at 24hpi. In column 1, infected worm is outlined (solid white line) with total number (n) of infected animals visualized in all replicates. Scale bar, 25μm. Columns 2, 3 & 4 show enlarged images of region boxed in column 1. Dotted lines indicate nuclei in G+AG (column 4) to enable visualization of nuclei in other columns. (B) Violin plot showing number of nuclei surrounded by genome smFISH signal in indicated strains. Kruskal-Wallis Multiple comparison test, *\*\*;p<0.01, ***; p<0.001, ****;p<0.0001.* (C) Quantification of percent infected animals exhibiting perinuclear localization of oRNA1 antigenome at 24hpi in indicated strains. Data are mean with standard deviation, N≥3, biological replicates. One-way ANOVA with Tukey’s multiple comparison test. *\*;p<0.05, ***;p<0.001.* (D) Representative northern blots showing viral siRNAs and U6 snRNAs detected using antigenome smFISH probes or U6 DNA oligonucleotide probes (end-labeled with ^32^P γ-ATP); 3μg of total RNA isolated from indicated strains at 72hpi were used for each blot N=3. (E) As in (D) using 2μg of total RNA from wildtype animals (72hpi) with or without terminator treatment. (N=3). % value indicates the increase or decrease (indicated by arrows) in viral siRNA intensity after normalization to U6 signal.

The 48 smFISH probes designed to bind antigenome, and the 27 that bind genome, were distributed along the length of the antigenome and genome, respectively, and by overlaying viral siRNA reads determined in a prior study (Ashe *et al*., 2013), we noted probes that should hybridize with positions of abundant viral siRNAs (**Supplemental Figures 4A-4D**); indeed, northern blot analyses using these probes detected small viral siRNAs (22 or 23 nts) in 72hpi wildtype animals but not in RNAi mutants (**Figure 4D**). We treated wildtype samples with “terminator”, which specifically degrades RNAs with 5’ monophosphates such as 1°, but not 2°, siRNAs. Upon quantification, we did not observe any significant difference in the levels of viral siRNA-sized products with terminator treatment (**Figure 4E**), indicating that the siRNAs detected by northern blot were predominantly 2° siRNAs at 72hpi. While it is difficult to compare the sensitivity of northern analyses with smFISH, these results suggest that 2° siRNAs from oRNA1 would be easier to detect by imaging than 1° siRNAs, which is not surprising given the abundance of 2° siRNAs (Ashe *et al*., 2013; Dahal *et al*., 2025). This combined with the lack of changes in the perinuclear signal in RNAi mutants that should affect 2° siRNAs, supports the idea that the perinuclear signal derives from antigenome or antisense 1° siRNAs.

### Perinuclear puncta do not colocalize with antiviral RNAi factors

Given the loss of perinuclear antigenomic signal in a subset of animals with deficiencies in the AVC, we considered the possibility that factors of the antiviral RNAi pathway (**Figure 5A**) might colocalize with the perinuclear antigenomic signal. We took advantage of existing strains for DCR-1, RDE-1 and DRH-3, a helicase related to DRH-1 that is involved in 2° siRNA amplification, that were tagged at the endogenous locus with 3XFLAG::GFP. We observed a diffuse distribution of DCR-1, RDE-1 and DRH-3 without infection (**Figures 5B, 5C and 5D**). As expected from prior studies (Phillips *et al*., 2012), in uninfected and infected animals we observed DRH-3 localized at perinuclear foci in the germline which served as an internal control (**Figure 5D**).

**Figure 5:**
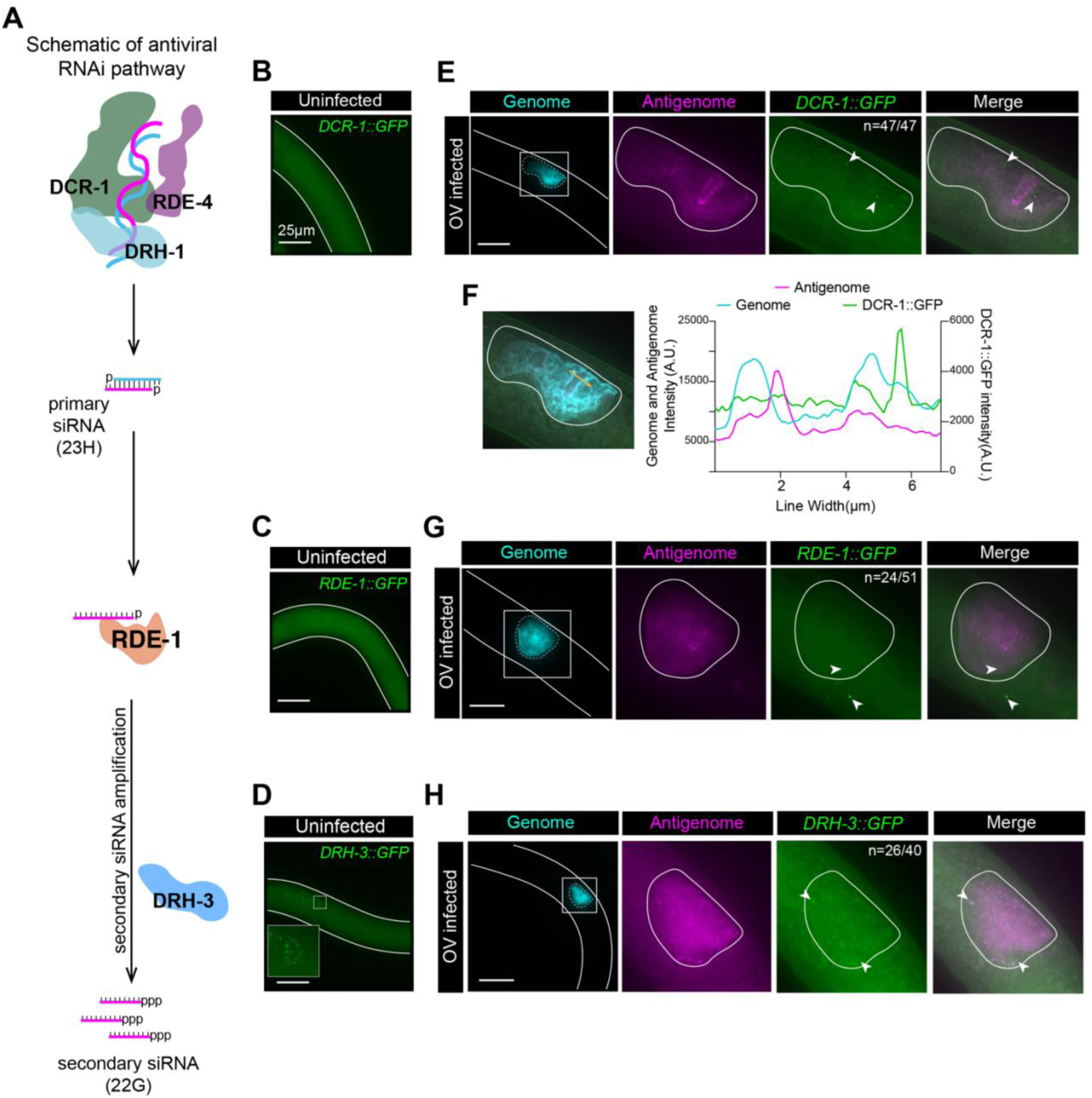
Perinuclear localized oRNA1 antigenomic signal does not colocalize with known antiviral RNAi factors. (A) Schematic of antiviral RNAi pathway. Maximum Intensity Projection images as labeled for DCR-1::3XFLAG::GFP, GFP::3XFLAG::RDE-1, DRH-3::GFP::3XFLAG, oRNA1 genome (cyan), oRNA1 antigenome (magenta) and merge in intestinal cells. Shown are localization patterns for uninfected animals (B, C, D) or at 24hpi (E, G, H). Boxed region in (D) shows an enlarged image of perinuclear localized DRH-3 in germline (inset). Columns 2,3 & 4 in E, G and H show infected cell (white solid line) in region boxed in column 1. White arrowheads point to GFP puncta. Number in column 3 (n) indicate the fraction of infected animals with GFP puncta in infected cells. (F) Line scan analysis of DCR-1::3XFLAG::GFP infected cell showing distribution pattern of genome, antigenome and DCR-1::3XFLAG::GFP. Yellow arrow shows path of scan. Over 40 infected animals were scored among 2 independent replicates for each strain. Scale bar, 25μm.

For all GFP-tagged proteins, DCR-1, RDE-1 and DRH-3, at 24hpi we observed distinct cytoplasmic puncta in cells with genomic signal (**Figures 5E, 5G and 5H**), and these puncta were clearly separated from perinuclear antigenomic puncta, as confirmed by line scan for DCR-1 (**Figures 5E and 5F**). However, a few cytoplasmic DCR-1, RDE-1 or DRH-3 puncta colocalized with cytoplasmic antigenomic puncta (**Figures 5E, 5G, and 5H**). Notably, DRH-1, a known interactor of DCR-1, also exhibits cytoplasmic punctate distribution in OV-infected cells (Batachari *et al*., 2024). The cytoplasmic punctate pattern for DCR-1 was observed in 100% of infected animals, and for RDE-1 and DRH-3, in around 50% of infected animals. Thus, although we observed that mutations in factors involved in the production of 1° siRNAs, but not those required for 2° siRNA amplification, showed reduced number of infected animals with perinuclear antigenomic signal (**Figure 4C**), at least for the factors tested, we do not observe the factors themselves localizing to the perinuclear region. Based on known mechanisms for antiviral RNAi, where 1° siRNAs would be expected to be proximal to the factors that generate them, these results are more consistent with the idea that perinuclear signal derives from the antigenome rather than 1° siRNAs; importantly, it is also possible that we are observing a previously unrecognized step of antiviral RNAi.

### dsRNA accumulates during OV infection and associates with oRdRP

Positive-strand viruses amplify their genome using the antigenome as template (Ball, 1994), and viral dsRNA arises during this process. dsRNA is often visualized using dsRNA-specific antibodies (de Faria *et al*., 2023), and to understand if dsRNA accumulation varies in wildtype and RNAi mutants, we used the J2 dsRNA-specific antibody with a dot blot assay, first showing that we could detect as low as 50pg of purified 800 basepair dsRNA (**Supplemental Figure 5**). We then collected total RNA from OV-infected (72hpi) and uninfected samples and again used the dot blot assay to monitor dsRNA accumulation. Control experiments showed that dsRNA was not detected without OV infection, and consistent with the fact that wildtype animals have efficient antiviral RNAi, we also did not observe dsRNA accumulation in wildtype animals (**Figure 6A**); however, given the limited sensitivity of this assay, we cannot rule out the existence of low levels of dsRNA in the wildtype background. Importantly, we detected dsRNA accumulation in all RNAi mutants tested (**Figure 6A**).

**Figure 6:**
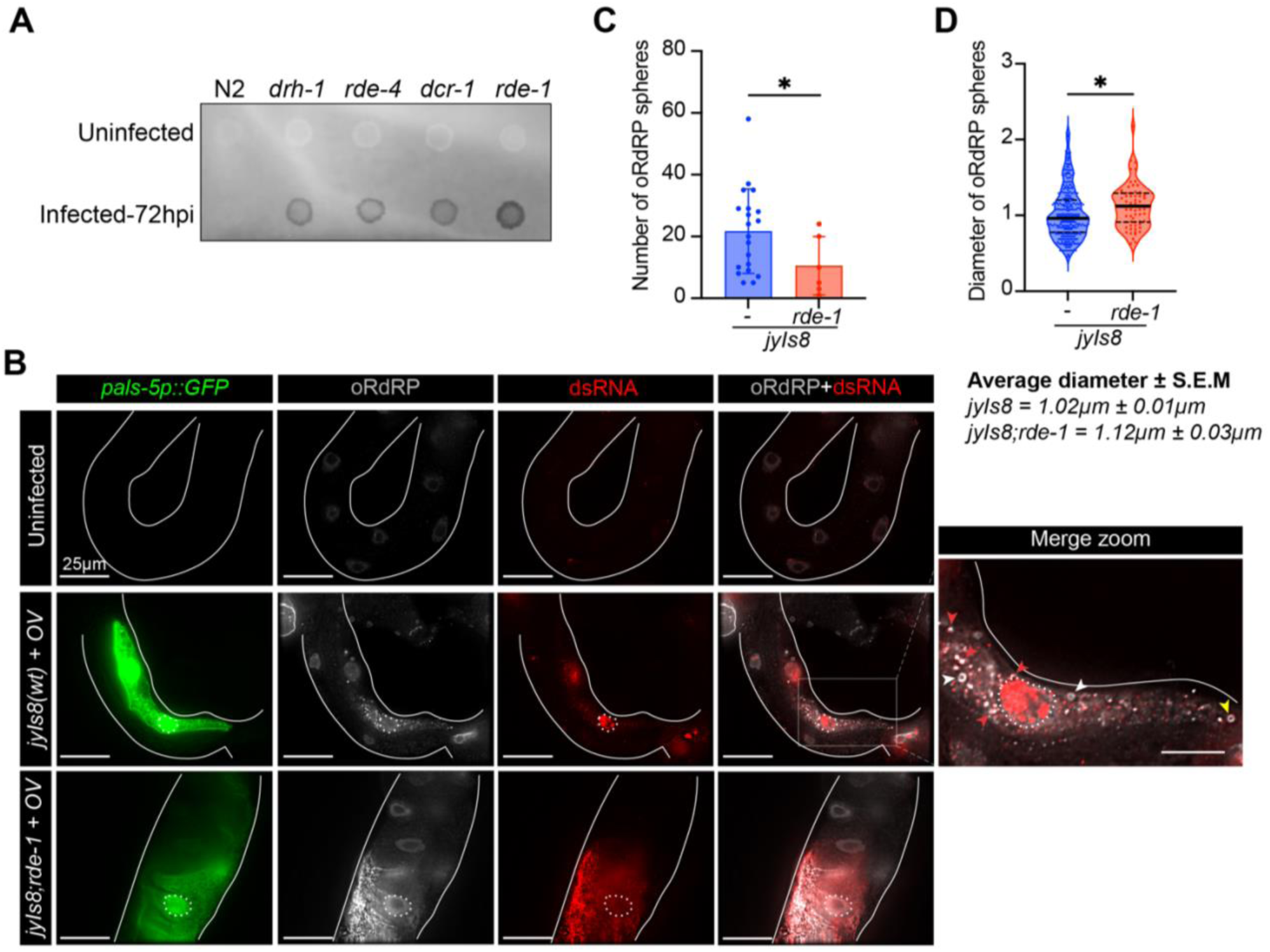
oRdRP is present in spherical structures or dsRNA-associated puncta in OV infected cells. (A) Representative dot blot spotted with total RNA from uninfected and OV infected animals at 72hpi; probed with J2 anti-dsRNA (J2) antibody. (N=3, biological replicates). (B) Representative images of uninfected or infected (24hpi) *jyIs8* (wt) or *jyIs8;rde-1* dissected intestines (white solid outline). Infection was signaled by expression of *pals-5p::GFP* (green) and also visualized were oRdRP (gray), dsRNA (red), merge (oRdRP and dsRNA). Nuclei (as marked by Hoechst) outlined with white dotted line. Red arrowheads, possible co-localization of oRdRP and dsRNA; white arrowheads, oRdRP spherical structures; yellow arrowhead, oRdRP sphere near dsRNA puncta. Scale bar, 25μm; merge zoom,10μm. N≥4 biological replicates with at least 5 animals per replicate for *jyIs8*; N=3 biological replicates with at least 5 animals per replicate for *jyIs8;rde-1*. Signal from oRdRP in intestinal nuclei indicates non-specific background staining. (C) Quantification of number of oRdRP spheres per infected animal in indicated strains. Unpaired t-test with Welch’s correction*; *, p=0.0424*. (D) Distribution of diameter of each sphere measured from the infected animals. Unpaired t-test with Welch’s correction*; *, p=0.0154*.

To evaluate whether the dsRNA signal was from viral or endogenous dsRNA, we looked for oRdRP colocalized with dsRNA in infected cells. We infected *jyIs8* (wildtype) or *jyIs8;rde-1* animals, picked GFP-positive infected animals and performed intestinal immunofluorescence assays using antibodies against oRdRP and dsRNA (J2). We observed that *pals-5p::GFP* expressing infected intestinal cells showed both oRdRP and dsRNA puncta, colocalized, in both *jyIs8* and *jyIs8;rde-1* animals (**Figure 6B; red arrowhead**). We also observed a strong nuclear dsRNA signal in oRdRP containing *pals-5p::GFP* expressing *jyIs8* intestinal cells that was not observed in uninfected animals or *jyIs8;rde-1* mutants (**Figure 6B, merge zoom**). The dsRNA that is colocalized with oRdRP is consistent with viral dsRNA being produced during replication, while the nuclear dsRNA more likely reflects endogenous intronic dsRNA induced in infected animals (Reich *et al*., 2018; Reich and Bass, 2019).

Interestingly, apart from oRdRP puncta, we noticed distinct oRdRP-containing spherical structures in the *pals-5p::GFP* positive intestinal cells (**Figure 6B, merge zoom, white arrowhead**). Over 75% of *pals-5p::GFP* positive wildtype animals exhibited at least one spherical oRdRP signal, with the rest only showing oRdRP puncta (*p=0.0046*); less than 50% of *rde-1* mutant animals exhibited oRdRP spheres. We quantified the total number of oRdRP spheres per animal and observed that wildtype animals showed between 5 to 58 spheres with an average diameter of 1.023μm (**Figures 6C and 6D)**. The number of spheres per animal was significantly lower in *jyIs8;rde-1* animals (*p=0.0424)*, with 0 to 25 spheres per animal and an average diameter of 1.122μm (*p=0.0154*), slightly higher than the observed diameter in wildtype animals. While we did not observe any dsRNA puncta inside the oRdRP spherical structures, we did find a few spheres closely associated with dsRNA puncta (**Figure 6B, merge zoom, yellow arrowhead**). Overall, we report previously uncharacterized lumen-containing spherical structures lined with oRdRP in wildtype background.

## Discussion

The abnormal morphology of the intestine in RNAi mutant *C. elegans* after infection with OV has been visualized with transmission electron microscopy (Félix *et al*., 2011), and the OV genome and oRdRP have been visualized using smFISH and immunofluorescence (Franz *et al*., 2014; Castiglioni *et al*., 2024). Here we add to this understanding by characterizing OV replication intermediates that include antigenomic sequences, dsRNA and oRdRP. Using quantitative, strand-specific methods, we demonstrate genome is in vast excess of antigenome (**Figure 1**), as found in other positive-strand RNA viruses (Novak and Kirkegaard, 1991; Félix *et al*., 2011; Shulla and Randall, 2015; Warncke and Knudsen, 2022; Genoyer *et al*., 2025). By utilizing a helicase mutant of DCR-1 (*dcr-1(mg375)*), we confirm the importance of helicase function to antiviral RNAi and efficient antiviral defense (**Figure 1; Supplemental Figure 1**), as suggested previously by in vitro experiments (Consalvo *et al*., 2024). Probes designed to hybridize with viral antigenome allow unambiguous localization of the antigenome after infection with most mammalian viruses (Shulla and Randall, 2015; Liu *et al*., 2019; Genoyer *et al*., 2025). However, in *C. elegans*, the existence of the antiviral RNAi pathway, which generates both sense and antisense siRNAs, precludes a straightforward interpretation of signal from hybridization to antigenomic probes. Here we take the first steps to address this issue, showing that denaturation protocols are required for detection of antigenomic, but not genomic, sequences in infected cells, and that antigenomic probes reveal both cytoplasmic and perinuclear signal, with the later altered in animals defective for 1°, but not 2°, siRNA pathways. We also report that oRdRP is present at the surface of lumen-containing spheres, and a decrease in the number of spheres in *rde-1* mutants suggests spheres may be related to host clearance of viral byproducts (**Figures 6B, 6C**). Our study offers an example of an invertebrate viral infection model and sets the stage for a more complete spatial and temporal understanding of the OV life cycle and host responses.

### What replication intermediates are detected with OV antigenomic probes?

Our studies establish the conditions necessary for detecting OV antigenomic sequences, but as yet, we cannot definitively assign the signal acquired with antigenomic probes as arising from antigenome or the antisense strand of an siRNA. That said, by comparison to known paradigms, in some cases we can choose the most parsimonious conclusion. For example, during infection with +ssRNA viruses, the antigenome is typically restricted to regions of viral replication in the cytoplasm (den Boon *et al*., 2024), and our observation of cytoplasmic antigenomic signal after OV infection is consistent with this. However, in addition to cytoplasmic puncta, using dsmFISH, we observed ∼80% of OV-infected wildtype animals with a perinuclear signal using antigenomic probes (**Figure 7A**), without the presence of other signatures of replication, such as oRdRP, genome, or dsRNA (**Figures 3A and 6B**). In the *C. elegans* germline, perinuclear regions are hubs for RNAi machinery (Phillips *et al*., 2012; Huang *et al*., 2024), but so far, such perinuclear granules have not been observed in somatic tissues, and we did not observe association of perinuclear antigenome with a subset of known RNAi factors (DCR-1, RDE-1, DRH-3; **Figures 5E, 5G, 5H**). Of course, there are other factors involved in steps in between RDE-1 loading and 2° siRNA amplification, such as RDE-8 (Tsai *et al*., 2015), and also in the subsequent targeting of the genome for degradation, which requires SAGO-2 (Seroussi *et al*., 2023), and it remains possible that these factors or others will be associated with the perinuclear antigenome.

**Figure 7:**
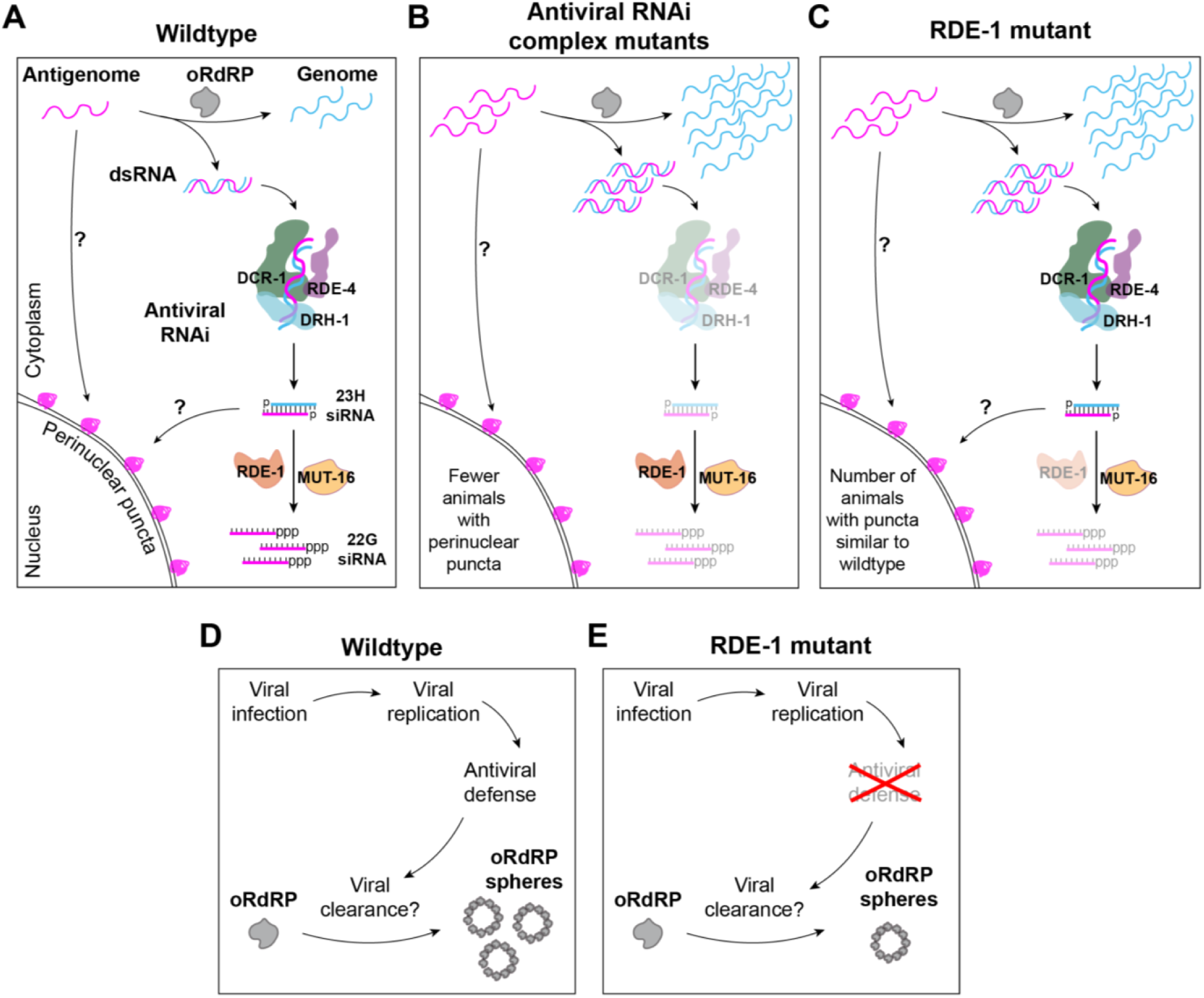
Models for antigenomic perinuclear signal and oRdRP spheres. (A-C) Cartoons showing observed replication intermediates and their potential intersection with the antiviral RNAi pathway. AVC factors are drawn to relative scale and adapted from Consalvo et. al. (Consalvo *et al*., 2024). A) In a wildtype animal, the perinuclear localized puncta could derive from OV antigenome or viral siRNAs. (B) In AVC mutants, single mutants of DCR-1, DRH-1, or RDE-4, we observed an increase in levels of viral RNAs and dsRNAs (Figures 1D, 1E, 6A and S1D-I), but the near absence of viral siRNAs (Ashe *et al*., 2013) leads to a reduction in the number of animals with perinuclear localized antigenomic signal. (C) Deficiencies in RDE-1 do not exhibit changes in the number of animals with perinuclear localized antigenomic puncta, despite these animals having an increase in viral RNAs and dsRNAs. (D) In wildtype animals with efficient antiviral defense, oRdRP spheres are abundantly formed as a part of viral clearance pathways. (E) In RDE-1 mutant animals, due to impaired antiviral defense, fewer oRdRP spheres are observed.

One of our most important observations in regard to the perinuclear antigenomic signal is that these puncta were dependent on the AVC factors required for producing 1° siRNAs (DCR-1, DRH-1, RDE-4) (**Figure 7B**) but are not affected by factors required for secondary siRNA biogenesis (RDE-1, MUT-16; **Figures 4A, 4C, 7C**). Regardless of whether the perinuclear signal results from 1° siRNAs or the OV antigenome, this indicates an important connection between OV infection and antiviral RNAi.

As evident from our strand-specific RT-qPCR assays, we observed a moderate increase in the antigenomic strand abundance in RNAi mutants relative to WT (**Figures 1D and 1E**). Thus, it is also possible that the loss of antigenomic perinuclear signal in primary RNAi mutants might correlate with an increased antigenomic signal in the cytoplasm, suggesting an impaired localization (**Figure 4A**). While images appear to suggest this, further quantification is necessary to conclude whether the increased cytoplasmic signal compensates for the signal from the perinuclear region in these mutants.

### What is the nature of dsRNA induced during OV infection?

During infection by both DNA and RNA viruses, dsRNA accumulates in infected cells, typically in the cytoplasm (Son *et al*., 2015). The large majority of existing studies detect dsRNA using commercially available antibodies (Corbet *et al*., 2022; de Faria *et al*., 2023), and the dsRNA is inferred to be of viral origin because it correlates with infection, and in some cases is observed associated with the viral RdRP (Andronov *et al*., 2024). Indeed, using an antibody to dsRNA, we and others have observed dsRNA accumulation in OV-infected intestinal cells (**Figure 6B**; (Batachari *et al*., 2024)). Previous observations of oRdRP show a cytoplasmic distribution in OV-infected cells (Franz *et al*., 2014), and in our studies, in wildtype and *rde-1* mutant animals, we find cytoplasmic dsRNA puncta associated with oRdRP, consistent with a viral origin (**Figure 6B**). These dsRNA-associated oRdRP puncta are possible viral replication hubs, similar to previous observations in other +ssRNA virus infections (Shulla and Randall, 2015; Andronov *et al*., 2024). We also observed closely associating genome and antigenome cytoplasmic puncta suggestive of replication hubs (**Figures 3F and 3H**). In future studies it will be important to determine whether antigenome can be observed to colocalize with oRdRP to confirm the presence of cytoplasmic replication hubs. dsRNA puncta that lack association with oRdRP could be partially processed dsRNAs or siRNAs as J2 can also recognize dsRNA as short as 14bp albeit with less affinity (Bou-Nader *et al*., 2025).

As in prior studies that used antibodies to detect dsRNA after viral infection of mammalian cells (Son *et al*., 2015), we also observed dsRNA in the nucleus. A robust nuclear J2 signal was observed only after viral infection (**Figure 6B, *jyIs8*, dsRNA**), in *pals-5p::GFP* positive cells, but not in neighboring *pals-5p::GFP* negative cells. While it is possible that this dsRNA is of viral origin, we favor the idea that it derives from infection-induced transcription of endogenous genes that contain intronic dsRNA, which are prevalent in many animals (Reich and Bass, 2019). Among the differentially-expressed genes produced during OV infection (Chen *et al*., 2017; Castiglioni *et al*., 2024), some do contain introns that encode dsRNA, such as *zip-1* and *uggt-2*, as defined in Reich et.al. (Reich *et al*., 2018). Interestingly the nuclear dsRNA signal is observed in wildtype-infected nuclei but not in *rde-1* animals (**Figure 6B**), possibly due to their inefficient antiviral response. This also highlights the importance of investigating antiviral responses in a wildtype background, since unique host responses to OV may not always be evident in mutant strains.

### What are oRdRP spheres?

Prior studies have only visualized oRdRP in RNAi mutant backgrounds at late larval stages, possibly explaining why these spherical structures were missed previously (Franz *et al*., 2014). While typical viral replication organelles are ∼300nm in diameter, we observed oRdRP-rimmed spheres (**Figures 6B**), that were larger, ∼1μm in diameter (**Figure 6D**), and in contrast to replication organelles, lacked association with dsRNA (**Figure 6B**). Further, even though *rde-1* mutant animals are more permissive to virus, they exhibited reduced oRdRP spheres (**Figures 6B and 6C**), consistent with the idea that spheres are not replication organelles. While higher resolution analyses are required to determine the nature and function of oRdRP sphere interaction with cellular organelles, one intriguing possibility is that these are structures associated with clearance, since viral RdRPs of +ssRNA viruses are not typically packaged into virions. Possibly, the oRdRP spheres we observe at 24hpi (**Figure 6B**) are formed from the remnants of replication sites targeted by the host for clearance after efficient antiviral defense (**Figure 7D**), which would be impaired in *rde-1* mutants (**Figure 7E**). In other viral infection models, viral RdRP is ubiquitinated and degraded through autophagy or proteosome degradation pathways (Liang *et al*., 2021), and between 12hpi and 24hpi with OV, genes related to ubiquitin-proteasome pathways are induced (Chen *et al*., 2017; Castiglioni *et al*., 2024). In *C. elegans*, infection with OV also induces GFP::LGG-1 puncta, which shows partial association with ubiquitin puncta (Bakowski *et al*., 2014). It will be important in future studies to determine whether or not the oRdRP in spheres is ubiquitinated by the host machinery and targeted for degradation through autophagy post viral replication.

## Materials and Methods

### *C. elegans* growth and maintenance

All *C. elegans* strains were cultured at 20°C under standard conditions unless specified (Brenner, 1974). The list of strains used in this study can be found in S1 Table. Some strains were provided by the CGC, which is funded by NIH Office of Research Infrastructure Programs (P40 OD010440).

### Orsay virus filtrate preparation

OV filtrate was prepared, and viral titer calculated, as described (Jiang *et al*., 2017). Briefly, JU1580 strain was cultured in S medium (T, 2006) and infected with OV for 3 days. Following infection, supernatant was collected and passed through a 0.22μm filter. The resulting filtrate was aliquoted and flash frozen at -80°C for long term storage. To calculate viral titer, *rde-1(ne219)* animals (1000 L1 per plate) were infected with serial dilutions (10-fold dilutions prepared in M9 buffer) of the viral filtrate.

### Orsay virus infection

#### For viral load, strand-specific PCR (ss-qPCR), northern blot and dot blot experiments

Strains were grown to gravid adults in liquid medium. Gravid adults were bleached (10% NaOH, 20% bleach), and embryos were synchronized in M9 buffer overnight at 20°C. For L3 infection experiments, 2000 synchronized L1s were plated in OP50-containing NGM plates and grown for 39 hours at 20° and infected with 0.1X OV for either 12 hours or 24 hours. For L1 infection experiments, 2000 synchronized L1s were plated in OP50-containing NGM plates and infected with 0.1X OV for 48 or 72 hours. The resulting infected and uninfected animals were washed in M9 buffer 4 times and collected in Trizol for RNA extraction.

#### For intestinal dissection

∼200 synchronized L1 animals were transferred to a plate containing OP50. Animals were incubated for 44-48 hours until they reached late larval stage (L3/L4). Animals were infected with OV (0.1X or 1X) for 24 hours and collected for dissections.

#### For smFISH experiments

∼300-500 synchronized L1s of respective strains were plated on OP50-containing plates and infected with OV (1X) at 20°C. Infected animals were collected after 24 hours of infection.

### RNA extraction, cDNA synthesis and qPCR

Total RNA from worms was collected using Trizol-chloroform-isopropanol extraction. Briefly, animals frozen in Trizol were thawed, and chloroform was added. Aqueous layer was separated, and RNA was precipitated using 100% isopropyl alcohol by incubating at room temperature for 30 minutes. The resulting pellet was washed and treated with DNase. DNase-treated RNA was then purified using Zymo RNA pure column as per manufacturer’s instructions. The resulting total RNA was used for all downstream analyses. cDNA was synthesized using Applied Biosystems cDNA synthesis kit as per manufacturer’s protocol for all experiments involving random hexamers. qPCR was performed using a QuantStudio3 instrument. Primers for qPCR are listed in S2 Table. Ct values were normalized to levels of *cdc-42*, and fold change was calculated using the ΔΔCt method (Livak and Schmittgen, 2001).

### Strand-specific RT-qPCR

We used strand-specific probes designed to bind the middle of one of the two Orsay RNA1 (oRNA1) strands; each probe also included a unique adapter sequence that did not bind within the viral genome or antigenome. To synthesize first strand cDNA, we incubated the probe with RNA and 10mM dNTP at 60°C for 10 mins, followed by reverse transcription at 37°C for 1 hour using the Applied Biosystems cDNA synthesis kit; enzyme was inactivated at 85°C for 20 minutes. We then used this strand-specific cDNA for qPCR. The unique adapter sequence was used as a forward primer combined with a reverse primer complementary to a region on the same strand of oRNA1, for qPCR. Primers are listed in S3 Table.

### Dot blot assay

Dot blot assays were performed as described (de Faria *et al*., 2023) with slight modifications. Briefly, 500ng of total RNA was diluted to 4μl and carefully spotted onto a supercharged nylon membrane. Once dried, the membrane was crosslinked under UV light (UV Stratalinker 1800, auto-crosslink setting). The cross-linked membrane was blocked in blocking buffer (1X PBS, 0.05% Tween-20, 50mg/ml sheared salmon sperm DNA, 5% skim milk) for 1 hour at room temperature (RT). The membrane was incubated with primary antibody (Anti-dsRNA:J2, 1:2000) overnight at 4°C. The membrane was then washed thrice with 1X PBST (1X PBS, 0.05% Tween-20) for 10 minutes at RT. The membrane was then incubated with secondary antibody (Anti-mouse IR dye 800CW, 1:20,000) for 1 hour at RT. The membrane was washed twice with 1XPBST for 10 minutes at RT, followed by two washes with 1X PBS for 10 minutes at RT. The membrane was then imaged using a LI-COR Odessey Infrared Imager. Antibodies were diluted in antibody dilution buffer (1X PBS, 0.05% tween-20, 2% skim milk). In-vitro synthesized and gel-purified ∼800bp dsRNA was used as control for the dot blot assay.

### Intestine dissection, immunostaining

Dissection and immunostaining protocols were based on (Batachari *et al*., 2024) with modifications as below. Briefly, GFP+ infected animals were picked onto 20μl of 5μM levamisole. Worms were dissected using sterilized surgical blades and transferred to 1X M9 buffer. Dissected intestines were washed twice with 1X M9, twice with 1X PBST and fixed in 4% paraformaldehyde at RT for 15 minutes. Fixed intestines were washed thrice with 1X PBST and blocked using a blocking solution (1X PBS, 0.5% Tween-20, 5% Bovine Serum Albumin, 0.5% Sodium azide) at 4°C overnight. Following blocking, intestines were incubated with Anti-oRdRP antibody (gift from Dr. David Wang (Franz *et al*., 2014)) and Anti-dsRNA (J2) antibody for 1-2 hours at RT. Unbound primary antibodies were washed off with 1X PBST, followed by incubation with secondary antibodies for 1-2 hours at RT. This was followed by two washes with 1X PBST and one wash with 1X PBST containing Hoechst (0.1ng/mL). For lipid droplet staining, LipidSpot^TM^ 610 Lipid droplet stain was diluted in 1X PBST and incubated for 30mins before Hoechst staining. Immunostained intestines were stored in 1X PBST at 4°C until mounting and imaging. Antibodies were diluted in blocking solution as per manufacturer’s recommendations. Antibody information is listed in S3 Table.

### smFISH

∼200 synchronized L1s of *jyIs8* or *jyIs8;rde-1* were plated on OP50-containing plates and infected with OV at 20°C. Infected animals were collected after 24 hours of infection. Animals were washed with 1X M9 buffer and 1X PBST (1X PBS, 0.05% Tween-20) and then fixed in 4% paraformaldehyde for 15 minutes at RT. Fixed animals were washed thrice with 1X PBST, followed by permeabilization in 70% ethanol in 1X PBST for at least 2 hours at 4°C. Permeabilized worms were washed thrice with 1X PBST to remove residual ethanol. If applicable, denaturation of dsRNA was performed by treating worms with 50mM sodium hydroxide for ∼30 seconds at RT (Genoyer *et al*., 2025) and immediately washing twice with 1X PBST. This was followed by a wash with 2X SSC buffer (Saline-Sodium Citrate), and then animals were pre-hybridized in 1 mL of Wash Buffer A (Stellaris® RNA FISH Wash Buffer A with nuclease-free water and 10% formamide). Samples were then hybridized to probes for the genome (27 probes) and antigenome (48 probes) (1μl of 12.5μM of each probe mix) in 100μl of hybridization buffer (Stellaris® RNA FISH Hybridization buffer with freshly added 10% formamide) per sample. Samples were incubated at 37°C overnight at 1200rpm (vortex). Post-hybridization, Wash Buffer A was directly added to samples followed by incubation at 37°C for 30 minutes (vortex). Pellet was washed with Wash Buffer A with Hoechst (0.1ng/mL) at 37’C or 30 minutes (vortex), followed by Stellaris® RNA FISH Wash Buffer B for 5 mins at RT on the nutator and then washed once with 1X PBST. Hybridized samples were stored in 1X PBST at 4°C until imaging. The sequences of the smFISH probes are listed in S4 Table.

### Imaging

Animals were mounted onto 2% agarose pads using Vectashield mounting solution. Images were captured using a DeltaVision Widefield Microscope with a 60X (oil) objective, Numerical Aperture (NA=1.42), equipped with a sCMOS digital camera using the Blue-Green-Orange-Far Red channels or Blue-Green-Red-Far Red channels. Multiple Z-slices spanning the entire thickness of the intestine with Z-stacks 0.2μM apart were captured. Images were captured and deconvolved using the softWoRx image processing software using standard settings. Individual channels were minimally adjusted in FIJI for better visualization. Unless otherwise specified, a single Z-slice was used to represent data. Acquisition setting for smFISH: Genome (Far-red), 5% transmission, 0.015-0.075 second exposure. Antigenome (Orange), 30-50%transmission, 0.200-0.500 second exposure. Note that antigenome channel was exposed for at least 10X longer than the genome channel.

### Quantification of perinuclear puncta distribution of oRNA1 antigenome

Deconvolved and processed images were opened on ImageJ, and colors were assigned for genome (cyan) and antigenome (magenta) channels. A composite image was created with a merge of these channels. Animals with at least one perinuclear puncta enriched for antigenome and negative for genome were considered a positive hit (visualized through line scan analysis). Randomization and blind quantification were performed with the help of Bass lab members to account for user bias.

### Quantification of oRdRP spheres

Deconvolved and processed images were used to perform oRdRP sphere diameter quantification on minimally adjusted Z-slices for the oRdRP channel. Progressive z-slices showing a point progressing to a circle and ending back in a point, was considered a sphere. Center z-slice of the sphere was determined, and the diameter (from a single z-slice) was calculated using the line tool on ImageJ. Randomization and blind quantification were performed with the help of Bass lab members to account for user bias.

### Heat shock inducible transgenic replicon system: heat shock induction protocol

Transgenic array, GFP positive (in pharynx) animals (4-5 L4s) were picked onto a fresh OP50-containing plate. Animals were incubated at 20°C for 3 days. Once they reached L4 larval stage, animals were heat shocked at 34°C for 2 hours. Animals were collected after 24 hours of recovery at 20°C for strand-specific RT-qPCR and northern blot assays.

### Standard curve for strand-specific RT-qPCR

In the region where the strand-specific primers were designed to bind oRNA1, a 500bp region was amplified using primers flanking a T7 promoter (genome) or SP6 promoter (antigenome). The purified PCR product was then used for in-vitro transcription to generate the genomic or antigenomic strand using T7 or SP6 RNA polymerase, respectively. Known numbers of purified ssRNA strands, ranging from 10^9^ to 10^2^ copies, were used for reverse transcription using strand-specific primers to generate cDNA. The resulting cDNA was used for qPCR to generate a standard curve with copy number on the x-axis and cycle threshold (Ct) values on the y-axis. Standard curve was generated, and the equation was obtained by fitting points to a semi-log line using GraphPad Prism software. The resulting equation was then used to determine copy number for strand-specific RT-qPCR analysis of infection experiments.

### Northern blot

Total RNA purified from early and late infected animals was heated at 65°C for 10 minutes in northern loading buffer containing ethidium bromide. Long transcripts (genome and antigenome) were resolved in agarose gel, while small RNAs were resolved in polyacrylamide-urea (17%(19:1), 8M Urea) gel. Buffers were prepared and northern blot was performed as described in (He, 2013). Membrane was then imaged under short UV light to capture ethidium bromide staining for the loading control. Membrane was pre-hybridized using hybridization buffer at 42°C for 20-30 minutes. 50μL purified, and end-labeled probes were mixed with hybridization buffer and added to the membrane for hybridization overnight at 42°C (37°C for siRNA probing); membranes were then washed with northern wash buffer thrice. Antigenome hybridized membranes were exposed to a PhosphorImager screen for 3 days; genome hybridized membranes were exposed for 2-3 hours; 72hpi sample containing membrane was only exposed for 15 minutes, viral siRNA probe-hybridized membranes were exposed overnight, U6 snRNA was visualized after 15minutes of exposure. PhosphorImage screens were imaged with a Typhoon Biomolecular Imager. The sequences of the northern blot probes are listed in S5 Table.

### CRISPR-Cas9 for *rde-4(uu71)* generation

*rde-4(uu71)* allele was generated with the CRISPR/Cas9 system as described (Reich *et al*., 2018).

## Abbreviations

RNA: Ribonucleic acid
DNA: Deoxyribonucleic acid
OV: Orsay Virus
RdRP: RNA-dependent RNA Polymerase
RNAi: RNA interference
dsRNA: double stranded RNA
siRNA: small interfering RNA
+ssRNA: positive, single strand RNA
OVIGs: OV induced genes
AVC: Antiviral complex
hpi: hours post infection
PCR: Polymerase Chain Reaction
RT-qPCR: Reverse transcription quantitative PCR
smFISH: single molecule fluorescence in-situ hybridization
dsmFISH: denature smFISH
GFP: Green Fluorescent Protein NGM Nematode Growth Medium
PBS: Phosphate buffer saline
RT: room temperature
ANOVA: analysis of variance

## Acknowledgements

We thank Bass lab members for helpful discussions and feedback. We thank Ryan Andrews for initial help with imaging of smFISH experiments. We thank Adam Hughes, Minna Roh-Johnson, Nidhi Raghuram, Ofer Rog and Sara Wong for helpful suggestions and feedback on microscopy data and methods. We acknowledge the Cell Imaging Core at the University of Utah for use of equipment (DeltaVision Ultra Widefield Microscope). Anti-RdRP (OV) antibody was a kind gift from Dr. David Wang. This work wias supported by funding to B.L.B. from the National Institute of General Medical Sciences (R35GNaM141262) and the National Cancer Institute of the National Institutes of Health (R01CA260414). B.L.B. is a Jon M. Huntsman Presidential Endowed Chair.

