## Supplemental material for "Characterization of Orsay virus replication intermediates in *Caenorhabditis elegans* reveals links to antiviral RNA interference"

**Supporting information**

**Supplemental Figure 1: Increase in viral load corresponds to increase in genomic strand in wildtype and RNAi mutants.**

Strand-specific RT-qPCR of purified genomic (A) or antigenomic (B) strands of oRNA1 were used to generate standard curves. Equations written in plot were used to determine copy number from strand-specific RT-qPCR shown in Figs 1 and S1. (C) Representative PhosphorImage of dot blot spotted with serial dilutions ($10^8$ to $10^2$ copies) of an ~500nt fragment of purified genome or antigenome RNA and probed using DNA oligonucleotides (end-labeled with $^{32}P$ $\gamma$-ATP) complementary to oRNA1 strands. Fold change (Genome/Antigenome) of wildtype and RNAi mutants of L3 infected (D) or L1 infected (E) animals at 12 or 24 hpi, as measured by strand-specific RT-qPCR. Data, mean with standard deviation. Representative northern blots of L3 (F) or L1 (G) animals for indicated times of infection points along with controls (JU1580, WUM32, WUM33) probed with DNA oligonucleotides (end-labeled with $^{32}P$ $\gamma$-ATP) specific to oRNA1 genome or antigenome. Ribosomal RNA (rRNA) as loading control was visualized with ethidium bromide after transfer to nylon membrane. Strand-agnostic Orsay viral load in L3 (H) or L1 (I) animals after 12 or 24 hpi. Data, mean with standard deviation. One-way ANOVA with Tukey's multiple comparison test; **, p-value<0.05; **, p-value<0.01*, N=3 . Related to Fig 1.

**Supplemental Figure 2: Perinuclear-localized antigenomic signal is not due to bleed-through from genomic channel.**

Representative images of infected *jyIs8;rde-1* animal (24hpi) probed with only antigenome probes after denaturation. Shown are signal from *pals-5p::GFP* (green), genome (cyan), antigenome (magenta), antigenome zoom alone or with the nucleus (Hoechst, blue). Scale bar 25μm. Related to Fig 3.

**Supplemental Figure 3: Establishing background signal in uninfected animals using genomic or antigenomic smFISH probes.**

Representative images of uninfected animals indicating minimal to no background signal of smFISH probes for oRNA1 genome (cyan), antigenome (magenta) and merge (genome and antigenome). Scale bar, 25μm. Related to Fig 4.

**Supplemental Figure 4: Comparison of locations of smFISH probes and viral siRNAs along the length of viral RNA.**

Primary siRNAs from 5'-dependent sequencing (A) or both primary and secondary siRNAs from 5' independent sequencing (B), from OV-infected wildtype animals were mapped to oRNA1 sequence (Data from Ashe et al., GSE41693). Viral siRNAs (antisense, magenta) across 20nt bins are plotted along the length of oRNA1 sequence on x-axis with read depth on the y-axis. Antigenomic smFISH probes are shown above with black bars. (C) and (D) are same as in A and B, respectively, but for sense viral siRNAs (steel blue) with genome smFISH probes shown below with dark blue bars.

**Supplemental Figure 5: J2 anti-dsRNA antibody can detect as low as 50pg of purified dsRNA.** Representative image of dot blot spotted with serial dilutions (50 ng to 5 pg) of in vitro synthesized and purified 800-bp dsRNA, probed with J2 anti-dsRNA antibody (N>3). Related to Fig 6.

**S1 Table:** List of *C. elegans* strains used in this study.

**S2 Table:** List of primers used in this study.

**S3 Table:** List of antibodies used in this study.

**S4 Table:** List of smFISH probes used in this study.

**S5 Table:** List of northern probes used in this study.

**Supplemental Figure 1**

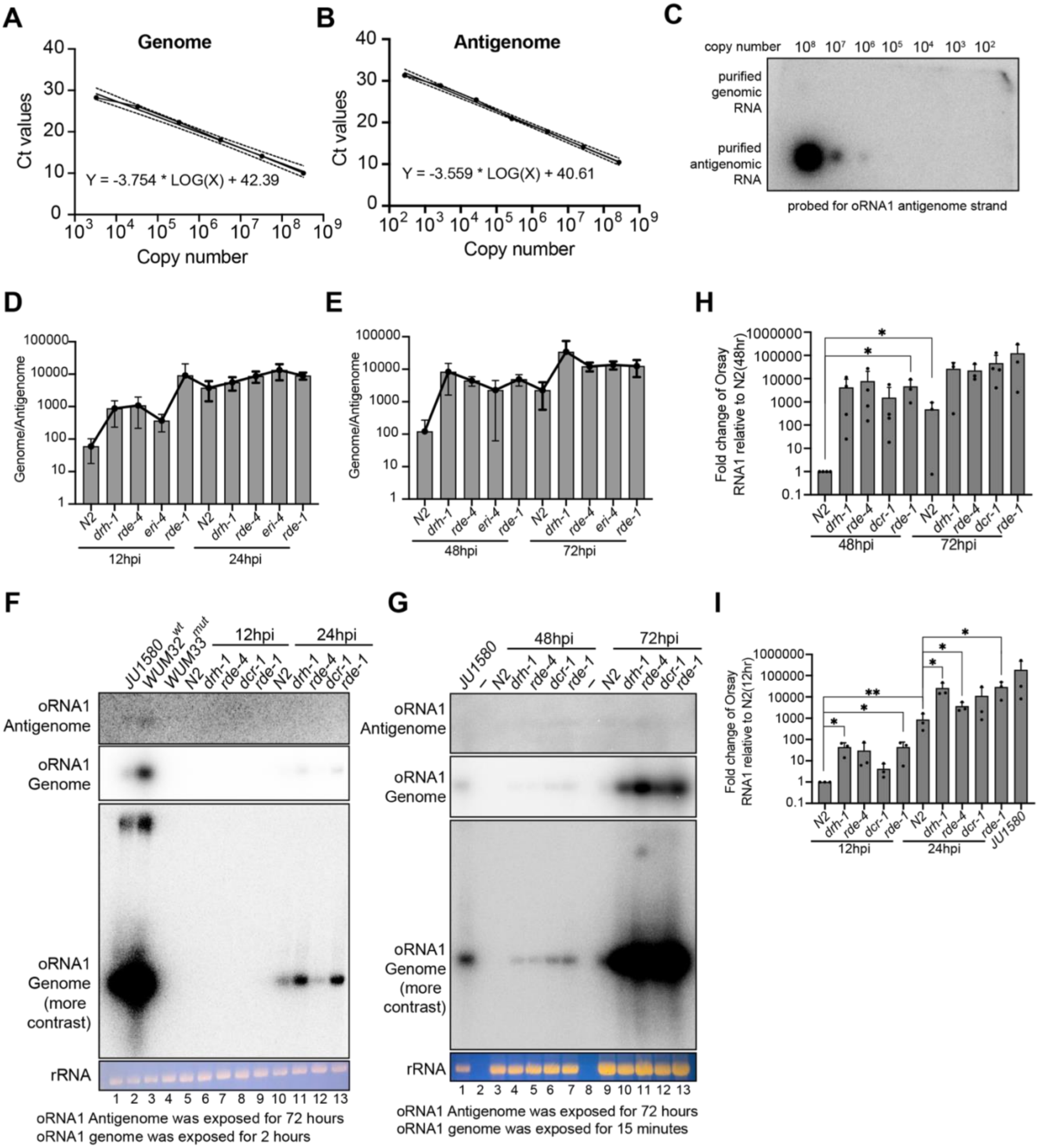

**Supplemental Figure 2**

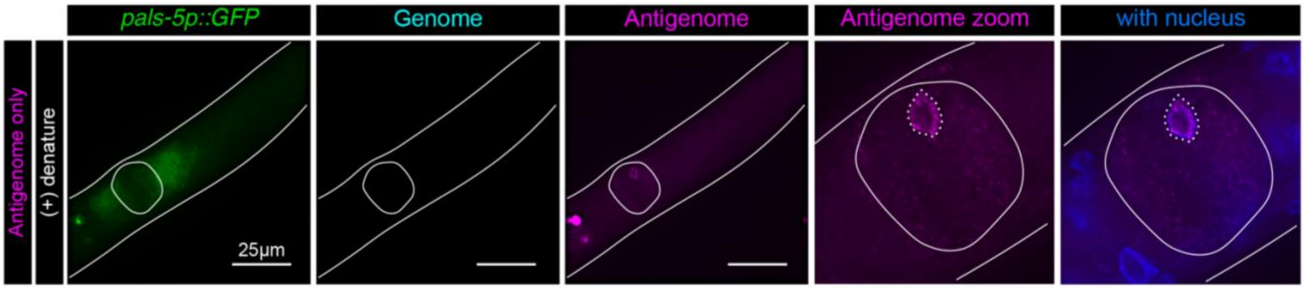

**Supplemental Figure 3**

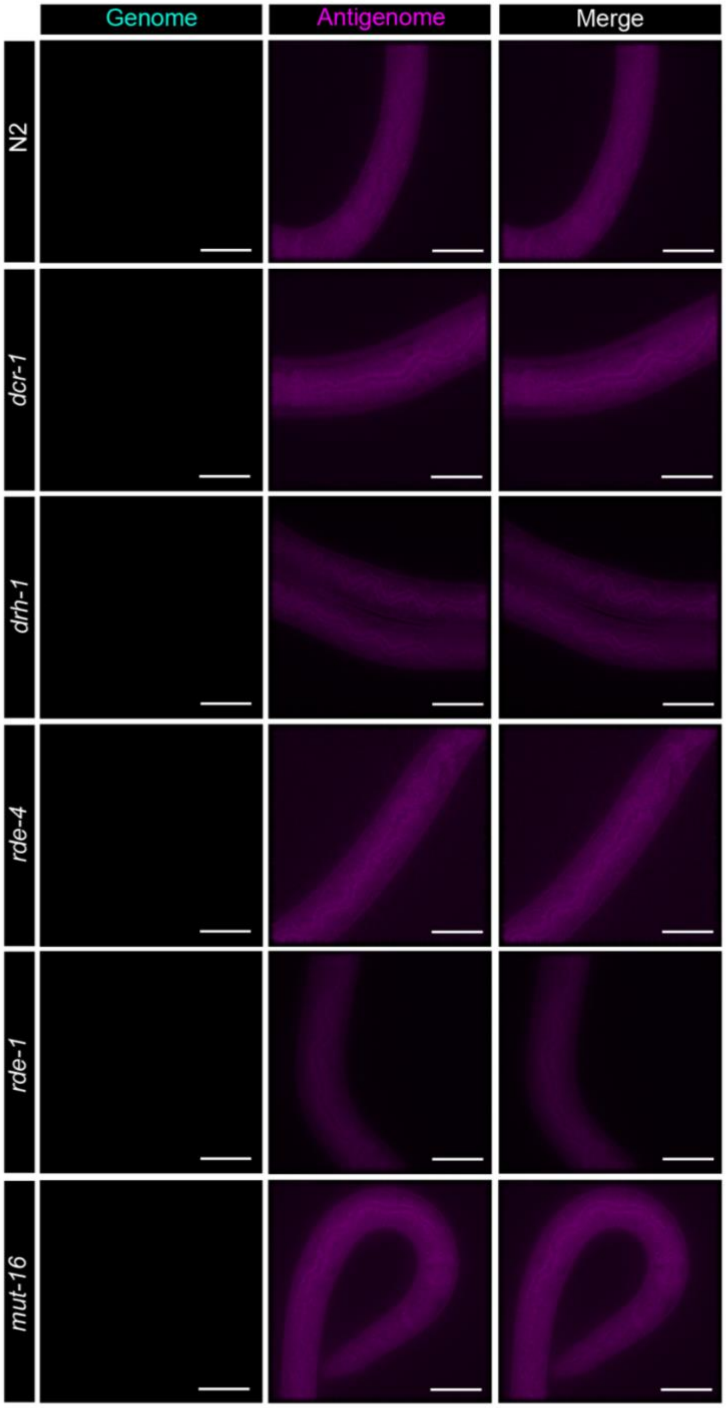

**Supplemental Figure 4**

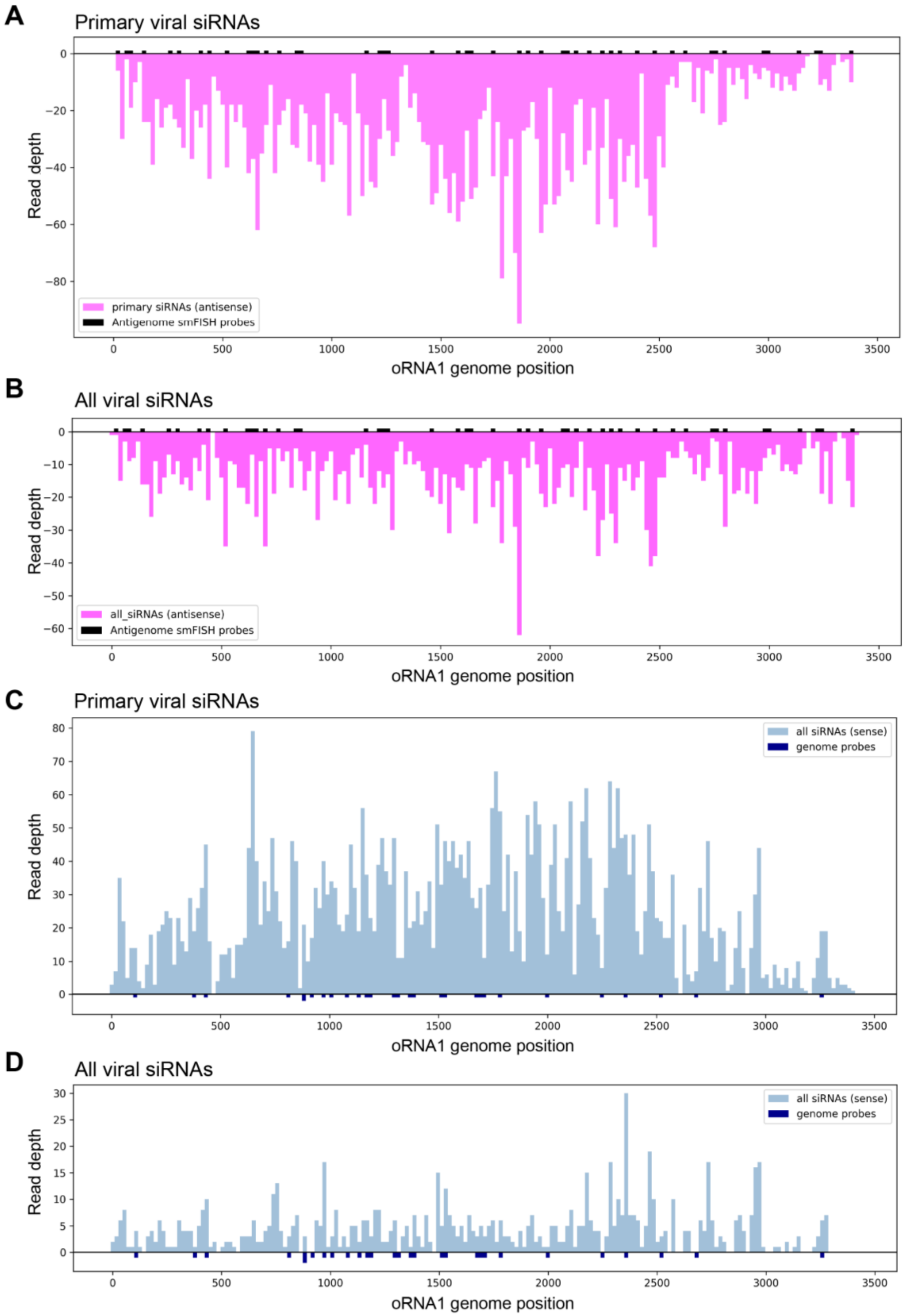

**Supplemental Figure 5**

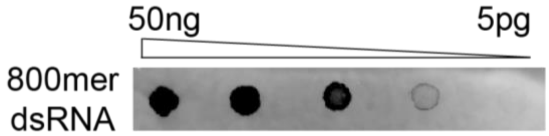

**S1 Table:** List of *C. elegans* strains used in this study.

| Strain | Genotype | Source (Citation) |
|---|---|---|
| N2 | +/+ (wildtype) | Caenorhabditis Genetics Center (CGC) |
| BLB256 | *drh-1(uu60) IV* | (Reich *et al.*, 2018) |
| BLB305 | *rde-4(uu71) III* | This study |
| YY70 | *eri-4(mg375) III* | [44] |
| WM27 | *rde-1(ne219) V* | CGC |
| JU1580 | *C. elegans* wild isolate with Orsay virus | (Félix *et al.*, 2011) |
| ERT54 | *jyIs8 (pals-5p::GFP; myo-2p::mCherry) X* | (Bakowski *et al.*, 2014) |
| BLB308 | *jyIs8 (pals-5p::GFP; myo-2p::mCherry) X; rde-1(ne219) V* | This study (virEx20 was outcrossed from WUM32). similar to WUM31 (Jiang *et al.*, 2017) |
| WUM32 | *jyIs8;rde-1(ne219); virEx20[PHIP::RNA1WT-1]* | (Jiang *et al.*, 2017) |
| WUM33 | *jyIs8;rde-1(ne219); virEx21[PHIP::RNA1D601A-1]* | (Jiang *et al.*, 2017) |
| NL1810 | *mut-16(pk710) I* | CGC |
| SHG1676 | *drh-3(ust352[drh-3::GFP::3XFLAG]) I* | CGC |
| SHG3112 | *dcr-1(ust624[dcr-1::GFP::3XFLAG]) III* | CGC |
| JMC217 | *rde-1(tor153[GFP::3XFLAG::rde-1]) V* | CGC |

**S2 Table:** List of primers used in this study.

| Name | Sequence | Comment |
|---|---|---|
| RNA1(GW194) | ACCTCACAACTGCCATCTACA | oRNA1 qPCR forward |
| RNA1(GW195) | GACGCTTCCAAGATTGGTATTGGT | oRNA1 qPCR reverse |

| | | |
|---|---|---|
| cdc-42-FP | AGCCATTCTGGCCGCTCTCG | qPCR forward |
| cdc-42-RP | GCAACCGCTTCTCGTTTGGC | qPCR reverse |
| RNA1-G-probe | CCAGATCGTTCGAGTCGT GACGCTTCCAAGATTGGTATTGGT | Probe for cDNA synthesis |
| RNA1-AG-probe | GCTAGCTTCAGCTAGGCATC ACTCAGTCAGCCTCAACAACC | Probe for cDNA synthesis |
| RNA1-G-FP | CCAGATCGTTCGAGTCGT | Strand-specific qPCR |
| RNA1-G-RP | ACCTCACAACTGCCATCTACA | Strand-specific qPCR |
| RNA1-AG-FP | GCTAGCTTCAGCTAGGCATC | Strand-specific qPCR |
| RNA1-AG-RP | GGTCTTGGTACGACGACTCG | Strand-specific qPCR |

**S3 Table:** List of antibodies used in this study.

| **Antibody** | **Host (dilution)** | **Obtained from** |
|---|---|---|
| dsRNA | Mouse Anti-dsRNA – J2 (1:200) | English Scientific and Consulting Kft. |
| oRdRP | Rabbit Anti-oRdRP (1:100) | David Wang's lab |
| Anti-Mouse | Donkey Anti-Mouse AF647 (1:1000) | Thermo Fischer Scientific |
| Anti-Rabbit | Goat Anti-Rabbit AF555 (1:500) | Thermo Fischer Scientific |
| Anti-Mouse | Goat Anti-Mouse (IR Dye 800 CW) | LI-COR |

**S4 Table:** List of smFISH probes used in this study.

| **Name** | **Sequence** | **Comment** |
|---|---|---|
| OV_RNA1_Sense_670_1 | CTAGATCCCAGTACAGTGGT | Genome probe |
| OV_RNA1_Sense_670_2 | CAAGGAGCAGTATCAGAACG | Genome probe |
| OV_RNA1_Sense_670_3 | TTAGACAGCCACAAGTTTCG | Genome probe |

| | | |
|---|---|---|
| OV_RNA1_Sense_670_4 | ATCACGATGTTGAGCGATTT | Genome probe |
| OV_RNA1_Sense_670_5 | GGGTCCATTCCACAACATAA | Genome probe |
| OV_RNA1_Sense_670_6 | GACATATGTGATGCCGAGAC | Genome probe |
| OV_RNA1_Sense_670_7 | TTGCACTGGAGAAAGTCATC | Genome probe |
| OV_RNA1_Sense_670_8 | CAGCCCTCATGACATAGTTG | Genome probe |
| OV_RNA1_Sense_670_9 | GGCGATGTAGACATATGTCG | Genome probe |
| OV_RNA1_Sense_670_10 | GTGGTCGTTCCAGTATGATG | Genome probe |
| OV_RNA1_Sense_670_11 | GTTCTTCGACGAGAGTGTTG | Genome probe |
| OV_RNA1_Sense_670_12 | CTTGTTCGGGTTAGTGACAA | Genome probe |
| OV_RNA1_Sense_670_13 | GTTCTCGATCACTTGCTCAG | Genome probe |
| OV_RNA1_Sense_670_14 | CTTGATCATAGCCTGGACAG | Genome probe |
| OV_RNA1_Sense_670_15 | CGATGGACAGATGGATCTTG | Genome probe |
| OV_RNA1_Sense_670_16 | CAAACTGGGAGATGTACCAC | Genome probe |
| OV_RNA1_Sense_670_17 | AGACGTTTGGCCATTAGTTT | Genome probe |
| OV_RNA1_Sense_670_18 | CTCTTGCCGTGAGTTTGATT | Genome probe |
| OV_RNA1_Sense_670_19 | CACCAACTCTTCGACGGA | Genome probe |
| OV_RNA1_Sense_670_20 | GTGGTCAAGATGATGGCGT | Genome probe |
| OV_RNA1_Sense_670_21 | AGGTTGGCGATCGTGTAT | Genome probe |
| OV_RNA1_Sense_670_22 | AGCGAGGTTGTGGTGGAGA | Genome probe |
| OV_RNA1_Sense_670_23 | GTTCTGATGGCCCTTCATC | Genome probe |
| OV_RNA1_Sense_670_24 | GATTTGCAGTGGCTTGCT | Genome probe |
| OV_RNA1_Sense_670_25 | ATGGTGCCTCGTGCAGAA | Genome probe |
| OV_RNA1_Sense_670_26 | CGAGGAAGGAGATGAAAG | Genome probe |
| OV_RNA1_Sense_670_27 | ATGGAGTCTGGGTTGTTGAG | Genome probe |
| OV_RNA1_AS_570_1 | GTCAACCTATGTGCCTAAAC | Antigenome probe |
| OV_RNA1_AS_570_2 | ATCTTCGGACGTTGCCAAAA | Antigenome probe |

| | | |
|---|---|---|
| OV_RNA1_AS_570_3 | AACAAGAACGATCCAGGTCC | Antigenome probe |
| OV_RNA1_AS_570_4 | GAGATTGGCAATTGGAGACC | Antigenome probe |
| OV_RNA1_AS_570_5 | CAATATCGAAGCCGTTGCAG | Antigenome probe |
| OV_RNA1_AS_570_6 | GGACGAAGTCGAGGAAACCG | Antigenome probe |
| OV_RNA1_AS_570_7 | ATGATGAACTCATGCGCTCA | Antigenome probe |
| OV_RNA1_AS_570_8 | ACTGCTGAGCAGTATTCGAA | Antigenome probe |
| OV_RNA1_AS_570_9 | TATGGACTGGAGAAACCGGT | Antigenome probe |
| OV_RNA1_AS_570_10 | GCTTTACCACTTGTTGTATT | Antigenome probe |
| OV_RNA1_AS_570_11 | GCGTTCCTCAACAAGATGAG | Antigenome probe |
| OV_RNA1_AS_570_12 | AGTTCAACGAGTCGTCGTAC | Antigenome probe |
| OV_RNA1_AS_570_13 | ATAGTAGCCGACATGGTATC | Antigenome probe |
| OV_RNA1_AS_570_14 | ATGTTGGAGACGACTCAGTC | Antigenome probe |
| OV_RNA1_AS_570_15 | CTCGAGGGACTAGACCAATA | Antigenome probe |
| OV_RNA1_AS_570_16 | AACACGATCACCAACGCTTT | Antigenome probe |
| OV_RNA1_AS_570_17 | TTACCATATCGCATCGGATG | Antigenome probe |
| OV_RNA1_AS_570_18 | ACTACCTCAACGAGATGCAT | Antigenome probe |
| OV_RNA1_AS_570_19 | CACAACTGCCATCTACATGA | Antigenome probe |
| OV_RNA1_AS_570_20 | AAAACACCCACTCCTTACTG | Antigenome probe |
| OV_RNA1_AS_570_21 | ACTCGGATTCTCGACATAGT | Antigenome probe |
| OV_RNA1_AS_570_22 | GAGTGGCTCAAAGCCAATAC | Antigenome probe |
| OV_RNA1_AS_570_23 | TCAATCTCGTCTACGGTACA | Antigenome probe |
| OV_RNA1_AS_570_24 | CTCGCAGAACAAGCAATCGG | Antigenome probe |
| OV_RNA1_AS_570_25 | TTGCTCTATTGGATCCAACG | Antigenome probe |
| OV_RNA1_AS_570_26 | AAAGTGTGGCTGTGCATGAG | Antigenome probe |
| OV_RNA1_AS_570_27 | TCGACAGTGTGCGGAACAAC | Antigenome probe |
| OV_RNA1_AS_570_28 | CATGCACCGTACAAAGCTAC | Antigenome probe |

| | | |
|---|---|---|
| OV_RNA1_AS_570_29 | GAACGTCACTAAGAAGGCCA | Antigenome probe |
| OV_RNA1_AS_570_30 | GTTTGGAACACCATCTACGA | Antigenome probe |
| OV_RNA1_AS_570_31 | AACGACCATCACACTCACAC | Antigenome probe |
| OV_RNA1_AS_570_32 | GAACATGCTGCTGATGGTAC | Antigenome probe |
| OV_RNA1_AS_570_33 | ACCCCTAAAACCAGGAAACA | Antigenome probe |
| OV_RNA1_AS_570_34 | TTCATGGTCGGATAACCATC | Antigenome probe |
| OV_RNA1_AS_570_35 | CAACCGAGAAGTGCTTGACG | Antigenome probe |
| OV_RNA1_AS_570_36 | TCGCTCTACTGTACATCATT | Antigenome probe |
| OV_RNA1_AS_570_37 | TACCCGTGTTCATCTTGAAA | Antigenome probe |
| OV_RNA1_AS_570_38 | TTGTCGGGCTACAACAACTG | Antigenome probe |
| OV_RNA1_AS_570_39 | TTGCTCGACATACTCTACGA | Antigenome probe |
| OV_RNA1_AS_570_40 | TCTCGGTGTCCAAATTCATG | Antigenome probe |
| OV_RNA1_AS_570_41 | TAAAACCCTCATGTATGCGC | Antigenome probe |
| OV_RNA1_AS_570_42 | TTTGATAGTAATCCTGTGCC | Antigenome probe |
| OV_RNA1_AS_570_43 | GACCGCTAAAGCGAATTGGA | Antigenome probe |
| OV_RNA1_AS_570_44 | ACTCAGGCTATCAAACAGCA | Antigenome probe |
| OV_RNA1_AS_570_45 | CTAAGTTCCACCTTGAAGTC | Antigenome probe |
| OV_RNA1_AS_570_46 | CACATTTGCTTGTGGTAGTG | Antigenome probe |
| OV_RNA1_AS_570_47 | CATGAGTAAACGGCAGCGTA | Antigenome probe |
| OV_RNA1_AS_570_48 | CGACTTGATACCGGCAATTG | Antigenome probe |

**S5 Table:** List of northern probes used in this study.

| Name | Sequence | Comment |
|---|---|---|
| OV_RNA1_Genome_1 | GATGGCCTCGACATATGTGATGCCGAGACA CGTTTCGAAC | Genome probe |

| | | |
|---|---|---|
| OV_RNA1_Genome_2 | GTGTGAGTGTGATGGTCGTTTGACCTCCAGAGTGGGC | Genome probe |
| OV_RNA1_Genome_3 | CACGTTTCGATGCACTGCGCGTTCTTCGACGAGAG | Genome probe |
| OV_RNA1_Genome_4 | CCGATAGTGAATGGCGCTCAGATCGTTCTGATGGCCC | Genome probe |
| OV_RNA1_Genome_5 | CTTGTTCTGCGAGCCTGTTCCGGGCGATTTGCAG | Genome probe |
| OV_RNA1_Genome_6 | CCGGCCGTGTACCATTTGGTATTGGCTTTGAGCCAC | Genome probe |
| OV_RNA1_Genome_7 | CTCGCGGATGGCATGCATCTCGTTGAGGTAGTCAGG | Genome probe |
| OV_RNA1_Genome_8 | GGCTTTCGCAATGGAGTCTGGGTTGTTGAGGCTGACTG | Genome probe |
| OV_RNA1_Antigenome_1 | CATCTTGACCACGGATAGGGCCACCAAGCGATATCAC | Antigenome probe |
| OV_RNA1_Antigenome_2 | CTTTGCCACCCACATGCCAACTATGTCATGAGGGCTG | Antigenome probe |
| OV_RNA1_Antigenome_3 | CGTCACTAAGAAGGCCAGCACGTCATACACGATCGCC | Antigenome probe |
| OV_RNA1_Antigenome_4 | GCGCGACGATCCGCATGCACCGTACAAAGCTAC | Antigenome probe |
| OV_RNA1_Antigenome_5 | GTTGCTCTATTGGATCCAACGCCGTTAACGCGCTGG | Antigenome probe |
| OV_RNA1_Antigenome_6 | CAAGCAATCGGATGGATGGACGCGGCATGGAAATGTG | Antigenome probe |

| | | |
|---|---|---|
| OV_RNA1_Antigenome_7 | CCAAGATGGACGCGGCAAAACACCCACTCCTTACTGC | Antigenome probe |
| OV_RNA1_Antigenome_8 | TCCATTGCGAAAGCCGGAGCCATGCTCGGATACAAAATAG | Antigenome probe |
| U6 snRNA probe | CTCTGTATTGTTCCAATTTTAGTATATGTTCTCGG | U6 probe1 |
| U6 snRNA probe | CACGAATTTGCGTGCATCCTT | U6 probe2 |
